# Perisomatic Features Enable Efficient and Dataset Wide Cell-Type Classifications Across Large-Scale Electron Microscopy Volumes

**DOI:** 10.1101/2022.07.20.499976

**Authors:** Leila Elabbady, Sharmishtaa Seshamani, Shang Mu, Gayathri Mahalingam, Casey Schneider-Mizell, Agnes L Bodor, J. Alexander Bae, Derrick Brittain, JoAnn Buchanan, Daniel J. Bumbarger, Manuel A. Castro, Sven Dorkenwald, Akhilesh Halageri, Zhen Jia, Chris Jordan, Dan Kapner, Nico Kemnitz, Sam Kinn, Kisuk Lee, Kai Li, Ran Lu, Thomas Macrina, Eric Mitchell, Shanka Subhra Mondal, Barak Nehoran, Sergiy Popovych, William Silversmith, Marc Takeno, Russel Torres, Nicholas L Turner, William Wong, Jingpeng Wu, Wenjing Yin, Szi-chieh Yu, The MICrONS Consortium, H. Sebastian Seung, R. Clay Reid, Nuno Maçarico Da Costa, Forrest Collman

## Abstract

Mammalian neocortex contains a highly diverse set of cell types. These types have been mapped systematically using a variety of molecular, electrophysiological and morphological approaches. Each modality offers new perspectives on the variation of biological processes underlying cell type specialization. Cellular scale electron microscopy (EM) provides dense ultrastructural examination and an unbiased perspective into the subcellular organization of brain cells, including their synaptic connectivity and nanometer scale morphology. It also presents a clear challenge for analysis to identify cell-types in data that contains tens of thousands of neurons, most of which have incomplete reconstructions. To address this challenge, we present the first systematic survey of the somatic region of all cells within a cubic millimeter of cortex using quantitative features obtained from EM. This analysis demonstrates a surprising sufficiency of the perisomatic region to identify cell-types, including types defined primarily based on their connectivity patterns. We then describe how this classification facilitates cell type specific connectivity characterization and locating cells with rare connectivity patterns in the dataset.

## Introduction

Electron microscopy volumes provide a unique perspective on neural circuits by enabling dense tracing of individual axons, dendrites and synaptic connections. Progress in recent years in data acquisition and dense segmentation have dramatically grown the capability to acquire large scale datasets.^1–7^ The size of these volumes allow for large numbers of cells to be analyzed and reconstructions of entire dendrites and local axons of mammalian neurons are now possible. However, it raises the challenge of accurately classifying tens or hundreds of thousands of cells. Doing so is necessary for many basic investigations, from co-registering neurons, to studying specific cell populations (including neuronal and non-neuronal cells), or being able to characterize the cell type specificity of connectivity at scale. Existing methods for automated cell-typing based on morphology often necessitate nearly complete axonal or dendritic reconstructions.^8–11^ Such reconstructions currently require manual correction to the segmentation, often referred to as proofreading, which is prohibitively time consuming at scale. Other definitions of cell-types require an understanding of the connectivity profile of individual axons, and therefore also require axonal proofreading. For example, a chandelier cell or basket cell is defined most clearly by the way they distribute their synapses onto its target neurons.^12–15^

In practice, when analyzing a large scale electron microscopy volume, one wants to intelligently invest proofreading efforts into the cells and cell types that are of interest to one’s study. Just as experimental systems require genetic tools to provide inexpensive access to rare cell populations that would otherwise be difficult to study with non-selective techniques, large scale electron microscopy requires computational tools to provide inexpensive access to specific cell types to facilitate further analyses. While the automated segmentation is very impressive in many respects, a significant amount of proofreading is required to clean and complete the reconstructions of cells. This means classifying, or even finding, cells based on specific output connectivity profiles is difficult in the dataset. Moreover, after proofreading, a single neuron reconstruction contains thousands of accurate synaptic targets to identify (Figure 1). A method that could identify cell-types in the dataset in a way that is insensitive to changes in proofreading and truncation is therefore of high utility, both to automate the classification of targets of proofread neurons and to help guide proofreading to cells that are more likely to have connectivity patterns of interest.

**Figure 1:**
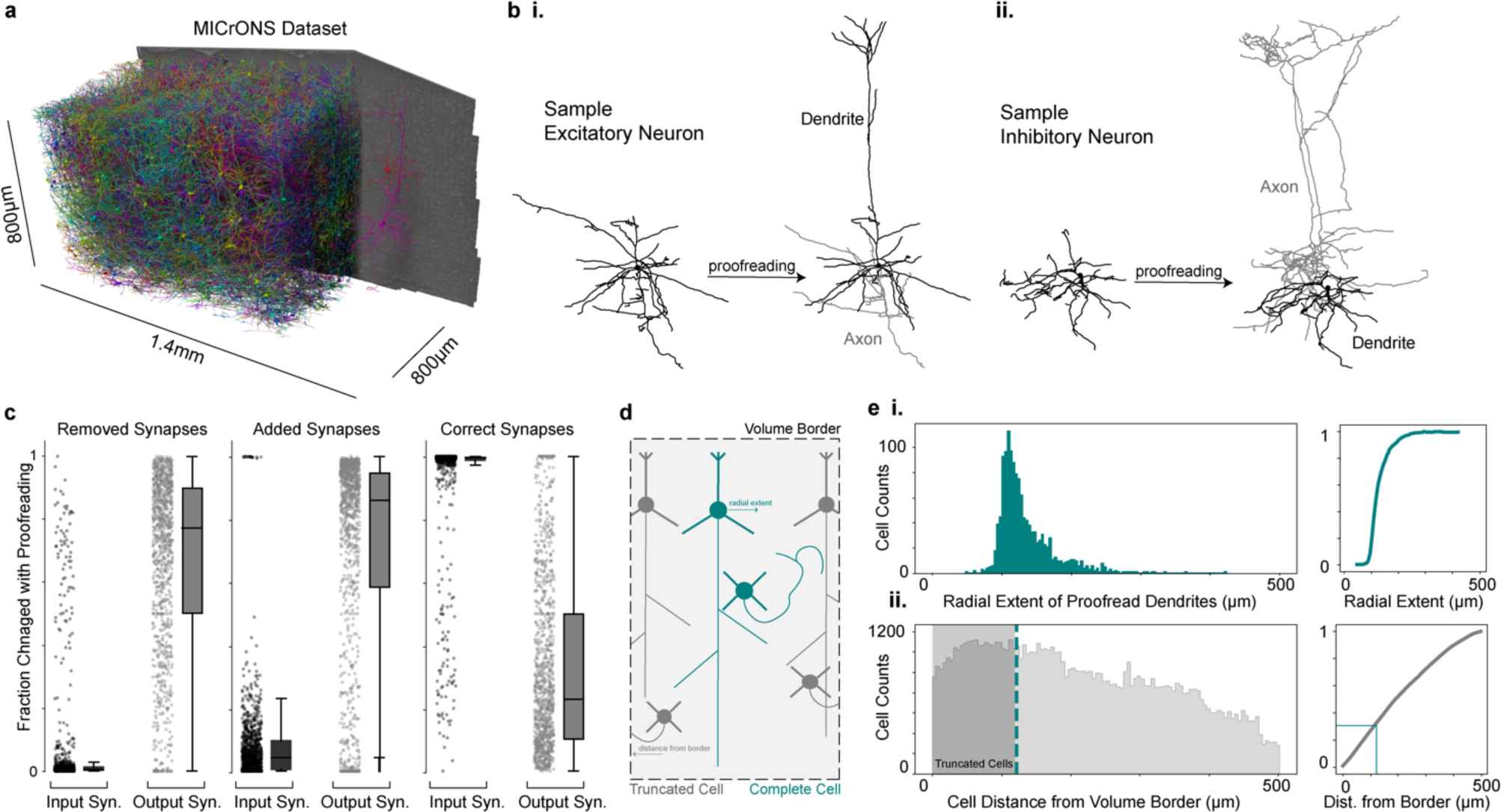
Large scale automated segmentations necessitate proofreading insensitive cell classifications. **a)** Rendering of a small fraction of neurons from the MICrONS dataset (1.4mm × 800μm × 800μm) covering all layers of cortex and multiple visual areas, with 1,207 rendered and then cutaway to reveal the full morphology of 2 selected neurons on the right portion of the dataset. **b)** Example neuronal morphologies before and after proofreading, **i)** excitatory neuron and **ii)** inhibitory neuron. **c)** Fraction of input and output synapses removed (left), added (middle) and maintained (right) after proofreading for 1,347 neurons. **d)** Neurons near the volume borders will inherently have truncated morphologies. **e) i)** Histogram of the radial extent of dendrites from a sample of 1,347 proofread neurons^16^ (left) and the cumulative distribution of those cells (right). **ii)** Histogram of the minimum distance from a volume border for all high quality nuclear detections (n=94,010) (left) and the cumulative distribution of those distances (right). The portion of cells which are less than the median radial extent (33% of cells) is indicated with teal shading and teal lines.

Here we describe a fast, scalable and computationally inexpensive approach which can address these problems. We first analyzed the somatic region of nearly 100,000 cells in the MICrONs dataset,^3^ a cellular compartment which contains morphological and connectivity based biological properties that, as we will present, differentiate cell-types. By analyzing only the somatic region of a cell, our analysis was generally robust to segmentation errors, unique per cell, and therefore insensitive to most proofreading changes. We included well-established features that are known to differentiate cells, such as somatic size and cortical depth, as well as novel features whose cell-type distinctions are less well recognized such as nuclear folding and soma synapse density. We further developed an unsupervised approach to describe the fine scale morphology of the perisomatic region of inhibitory cells, and demonstrate that it varies across major inhibitory subclasses. With these features in hand, we address the need for dataset wide cell-type labels outlined above, by training a hierarchical classifier to identify basic cell classes across the entire dataset. We demonstrate the utility of perisomatic features to facilitate the targeted search for rare cell types across a dataset. This method is already being used to reveal fundamental aspects of cell type specific wiring of mammalian cortex.^3,16–18^ More broadly, the efficacy of this approach provides a roadmap for how to develop a scalable platform for leveraging local features of cells to infer cell-type classifications across large scale image data.

## Results

### Segmentation Quality Varies Across Neuronal Arbors

We analyzed the larger portion of the MICrONS dataset, a 1.4mm × 800μm × 800μm volumetric serial section EM dataset from mouse visual cortex,^3^ that contains a dense segmentation of cells along with a nucleus segmentation and large scale synapse detection (Fig. 1a).^2,19^ This dataset includes 94,010 high quality nuclear detections fully enclosed within the boundaries of the volume (see methods) and spans cortical Layer 1 through to the white matter. For most cells, high quality cellular segmentation requires proofreading to clean and complete the reconstructions, particularly axons (Fig. 1b-c). Most false mergers are of axonal fragments, leading most outputs of unproofread axons to be incorrect (Fig. 1c). When axonal proofreading is invested in an individual cell, it creates an elaborate object to analyze with thousands of postsynaptic targets. In order to analyze the cell-type specific connectivity pattern of that single reconstructed cell (examples in Fig. 5-6), each of those post-synaptic cells requires a cell type label. Dendrites on the other hand are quite precise, as 75-95% of the 1,000 to 15,000 synapses detected on reconstructed axons can be mapped to their soma in the MICrONs dataset with more than 99% accuracy (Fig. 1c). However, many of these targets have incomplete reconstructions themselves because even for mm^3^ scale volumes about a third the cells are close enough to the edge to have their dendrites be truncated (Fig. 1d-e). This level of truncation across cells, whether due to segmentation errors or proximity to the volume border, led us to investigate alternative methods for cell-typing that would be insensitive to a cell’s dendritic and axonal reconstruction status.

### Perisomatic Features Vary Across Cortical Cell Classes and Subclasses

The somatic region of the cell is an attractive location for such a method to focus on because it is a unique region per cell and its limited spatial extent means it is rarely truncated. The automated reconstruction of the somatic region is typically precise and complete, with few changes during proofreading, if any (Fig. 2a). Moreover, the soma also has unique biological processes occurring within it, which led us to investigate if information within the perisomatic region could enable cell classification.

**Figure 2:**
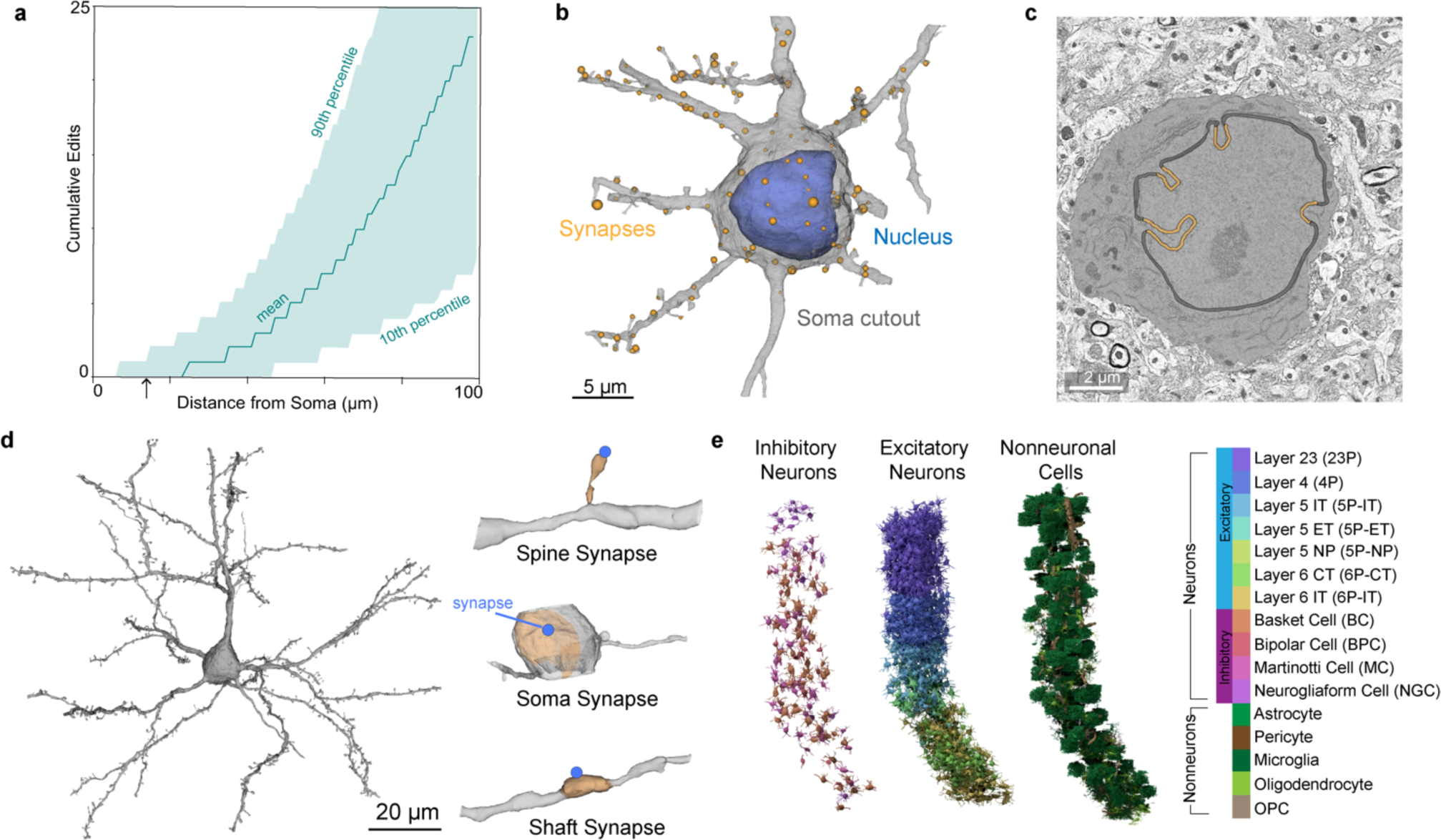
Perisomatic region of cortical cells. **a)** A measure of the distance from the soma for each edit that was made to the segmentation during proofreading of 1,347 cells. Average noted by the teal line, area chart notes the 10th and 90th percentile across all cells. Arrow notes 15 microns. **b)** Example cell demonstrating the extent of mesh information used to extract somatic, nuclear and synapse features. All cell meshes were restricted to 15 microns from the center of the nucleus. **c)** Representative example of nuclear infolding in a single electron microscopy image. The soma is highlighted in gray, black represents the nuclear envelope and orange are the areas classified as infolded based on the shrink wrap method (see Methods). **d)** Example cell demonstrating the extent of mesh information used to extract post-synaptic features (left) and three example post-synaptic shapes (PSS)(right). All synapses included in the PSS analyses were within 60 microns from the center of the nucleus. **e)** A 3D rendering of the somatic cutouts from all the cells from a 100 micron column that was densely reconstructed for which manual labels were given. Cells rendered are organized by their cell class and colored by their cell subclass according to the color scheme displayed.

We began by systematically extracting geometric properties of the nucleus and soma within 15 microns from the center of the nucleus (Fig. 2b). For nuclei, this included volume, surface area, and depth from the pial surface. The 3d nuclear segmentations provide a detailed view of an interesting feature of neuronal nuclei, their tendency to form infoldings of their membranes, sometimes also referred to as invaginations. We quantified the fraction of nucleus membrane area that was within an infolding using a geometrical method that shrink wraps the nucleus and labels membrane at least 150 nm from that wrap as part of an infolding (Fig. 2c) (see methods). The nucleus of different cell types has been described as having different degrees of infolding,^20,21^ but a systematic quantification has not been done across cortical types. We also calculated similar geometric properties of the somatic region of cells (see methods): the total volume, surface area, the ratio of the nucleus volume to the soma volume, and the distance from the centroid of the nucleus to the centroid of the soma. To capture information about neuronal connectivity, we also measured the number and surface density of synaptic inputs detected on the somatic region of the cell. Together these somatic and nucleus features represent a feature space that was extracted for most cells (75% of nuclei detections, see Methods). For a subset of neurons, we also analyzed the nano-scale structure of the postsynaptic compartments, what we are terming a “post synaptic shape” (PSS) (see methods) within 60µm of the nucleus center (Fig. 2d).

In order to develop a dataset wide cell-type classifier, we used a densely reconstructed and manually annotated column of 1,619 cells across all layers of primary visual cortex (Fig. 2e).^3,16^ This column included excitatory neurons (1,115), inhibitory neurons (143), and non-neurons (361) with expert annotated labels for cellular classes and neuronal subclasses (Excitatory: Layer 2/3, Layer 4, Layer 5 inter-telencephalic (IT), near-projecting (NP) and extra-telencephalic (ET), Layer 6 IT (IT) and corticothalamic (CT) and Inhibitory: Martinotti/non-Martinotti cell (MC), Basket cell (BC), Bi-polar cell (BPC) and neurogliaform cell (NGC), Non-neurons: astrocyte, oligodendrocyte precursor cell (OPC), oligodendrocyte, microglia, pericyte) (Fig. 2e). The cells in this column served as the ground truth throughout the rest of our analyses (see Methods). Though, it should be noted that our approach easily incorporates alternative labels as cell-type definitions evolve.

How effective are different features alone in separating cells at different levels of classification within the cortex? To answer this question we plotted various individual features, and trained classifiers to distinguish cells at different levels of granularity. Nucleus features alone were nearly sufficient to separate neurons from non-neuronal cells (cross-validated classification accuracy 90%, Extended Data Table 1). Non-neuronal cells had generally smaller nuclei compared to neuronal cells, though astrocytes overlap in this distribution with the smallest neurons. Each non-neuronal cell class exhibited a distinct range and consistency in their nucleus volume across the layers of cortex (Fig. 3a.i). A nucleus-only classifier was able to identify non-neuronal subclasses with a cross-validated accuracy of 94% (Extended Data Table 1). Nucleus features of excitatory neurons across the dataset recapitulated expected laminar organization, wherein the borders between layer 2/3 (L23), layer 4 (L4), layer 5 (L5), and layer 6 (L6) are all demarcated by shifts in the distribution of nucleus volumes. (Fig. 3a.i). Inhibitory cells, on the other hand, had less striking laminar patterns, but with more overall variation. They had a wider variation of nucleus volumes, which overlapped with those of excitatory cells, with the exception of the larger layer 5 excitatory neurons (Extended Data Fig. 1).

**Figure 3:**
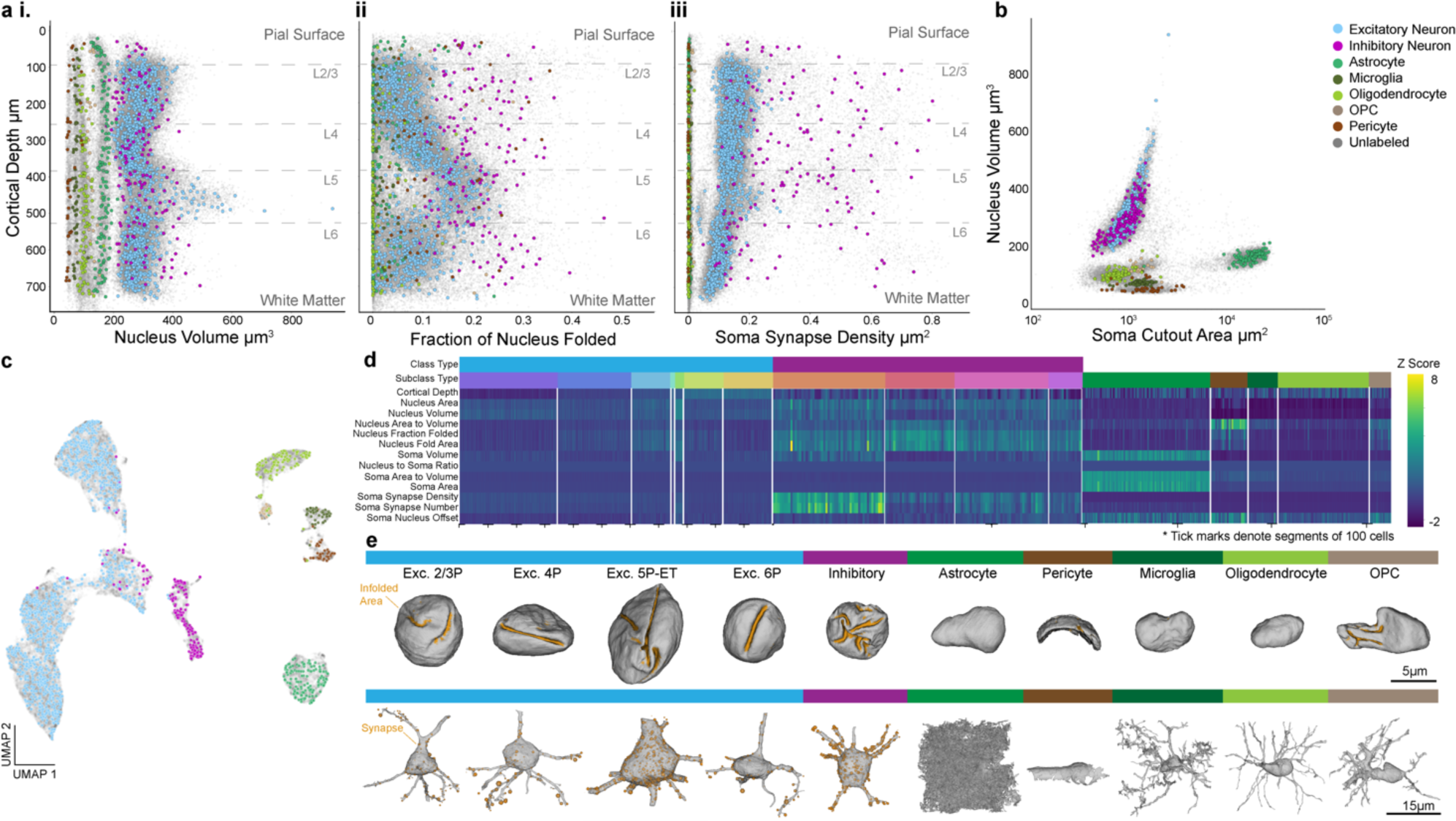
Variations of nucleus and somatic features show stark laminar and cell-class based distinctions. **a) i)** Nuclear volume μm^3^ **ii)** fraction of nuclear membrane infolded and **iii)** somatic synapse density μm^2^ plotted against distance from the pial surface in microns. Cortical layer boundaries are noted by the dashed lines. **b)** Somatic surface cutout area in μm^2^ (within 15µm from the nuclear center) plotted against nuclear volume μm^3^. **c)** 2D UMAP embedding of all neuronal and nonneuronal cells inferred from somatic features, nuclear features and cortical depth. **d)** Z-scored feature matrix representing all the somatic and nuclear features on the manually labeled cells from the cortical column. Cells are organized by their annotated subclass. Dashed marks along the × axis denote segments of 100 cells (1115 excitatory neurons, 143 inhibitory neurons, 361 nonneurons). For all plots, manually labeled cell classes are represented in color (1,619) and unlabeled examples in light gray (n=92,391). **e)** 3D mesh renderings of representative examples of different neuronal and non-neuronal cell classes. In the top row, nuclei displayed with the folded surface area highlighted in orange. Corresponding cell bodies displayed in the bottom row with somatic synapses in orange. Sphere size corresponds to predicted synapse size from the synapse detection model.^3^

The fraction of membrane inside an infolding also varied widely and systematically depending on depth (Fig. 3a.ii, 3e). Layer 2/3 neurons had largely smooth nuclear membranes. There was a clear gradient of infolding within layer 4. All layer 5 excitatory cells had high degrees of infolding, despite the notable diversity of cell types and sizes within that population, which was reflected in the increased variation of nucleus volume in that layer.^22^ Infolding dropped sharply again in layer 6 (Fig. 3a.ii). On the other hand, inhibitory nuclei had 15-30% of their membrane within an infolding, regardless of their position within cortex. This made them quite distinct from excitatory neurons in layer 1, 2/3, 4 and 6 of cortex, but highly similar to those in layer 5 (Fig. 3a.ii, Extended Data Fig. 1). Non-neuronal cells generally did not have infoldings, though microglia, OPCs and oligodendrocytes had less spherical and convex shapes (Fig. 3e). Pericytes had the smallest overall volumes with shapes dominated by their close apposition to the vascular walls (Fig. 3e).

Two features alone, nucleus volume and soma cutout area, revealed a surprisingly striking separation between the major cell classes found in the brain (Fig. 3b). In particular, neurons were separated from all non-neuronal classes and microglia, oligodendrocytes, OPCs, astrocytes, and pericytes occupy largely distinct portions of this 2-dimensional space. The large surface area measurement for astrocytes was explained by the high density of their processes near the soma. Moreover, the high prevalence of segmentation mergers of pericytes with cortical vasculature resulted in variability in their soma size features as represented by the range in soma cutout area (Fig. 3b). Including the somatic features along with the nucleus features, we trained a classifier to distinguish neurons, non-neurons and erroneously segmentations from each other with a cross validated accuracy of 95.6%, and a classifier on non-neuronal classes with 97.5% accuracy (Extended Data Table 1).

Excitatory neurons showed a consistent synapse density that varied slightly in a linear fashion with depth through the cortical volume. There was a notable increase in variation in layer 5 that correlated with the three subclasses found there with ET cells having larger synapse densities, NP cells with low synapse densities and IT cells in between (Fig. 3a.iii., 3e). Inhibitory neurons exhibited less laminar variation in somatic size. Inhibitory cells had much larger density of somatic innervation than excitatory cells, but also have a much wider degree of variation, reflecting previously recorded diversity of inhibitory subclasses (Fig. 3a.iii, 3e).^19,23–26^ Unsurprisingly, all nonneuronal cells have many fewer somatic synapse counts and thus are clearly distinct from neurons across laminar layers (Fig. 3a.iii). Classifier accuracy for excitatory subclasses was high (90% Extended Data Table 1), with most confusion surrounding IT cells located at laminar borders, general locations expert annotators can reasonably disagree. Notably, accuracy was high across the layer 5 cell types (99% for NP, 85% for IT, and 87% for ET).

To gain a broader understanding of the perisomatic feature landscape, we computed a low-dimensional embedding based on both nucleus and somatic features (Fig. 3c). Consistent with the diversity observed in individual features, low dimensional embeddings of the feature space across all the cells in the dataset (gray, n = 92,391) reflected the variations observed from the manually labeled cortical column (cell class colors, n = 1,619). Non-neuronal cell classes occupied distinct areas of the feature space whereas excitatory neurons were primarily organized by cortical layers (Extended Data Fig. 1). Inhibitory neurons were largely restricted to a distinct cluster within this space, with some cells overlapping with cortical layer 5 cells likely due to the increase in nuclear infolding in those excitatory neurons. Although there were broad average differences in the nucleus and somatic features between the major interneuron subclasses (Fig. 3d), our attempts at building classifiers based on those features produced some confusion (accuracy of 90%) (Extended Data Table 1).

### Proximal Dendrites of Inhibitory Neurons Vary in Distributions of Post-Synaptic Morphologies

At the same time, it was clear that the ultrastructure of inhibitory neuron peri-somatic regions was diverse in ways that the soma and nucleus features did not capture. Upon inspection, proximal inhibitory branches varied in caliber and surface texture, from smooth and uniform to covered in small spine-like protrusions (Fig. 4a). In order to take advantage of this information, we developed a method to summarize these fine shape statistics of the proximal arbor.^27^ Since automated spine detection is a challenging technical problem, we instead use automated synapse detections to identify areas on the dendrites where changes in fine structure (e.g. spines) may occur. This approach gave us the added advantage of combining information about synaptic innervation with the fine morphological structure of the postsynaptic neuron surrounding any given synapse.

**Figure 4:**
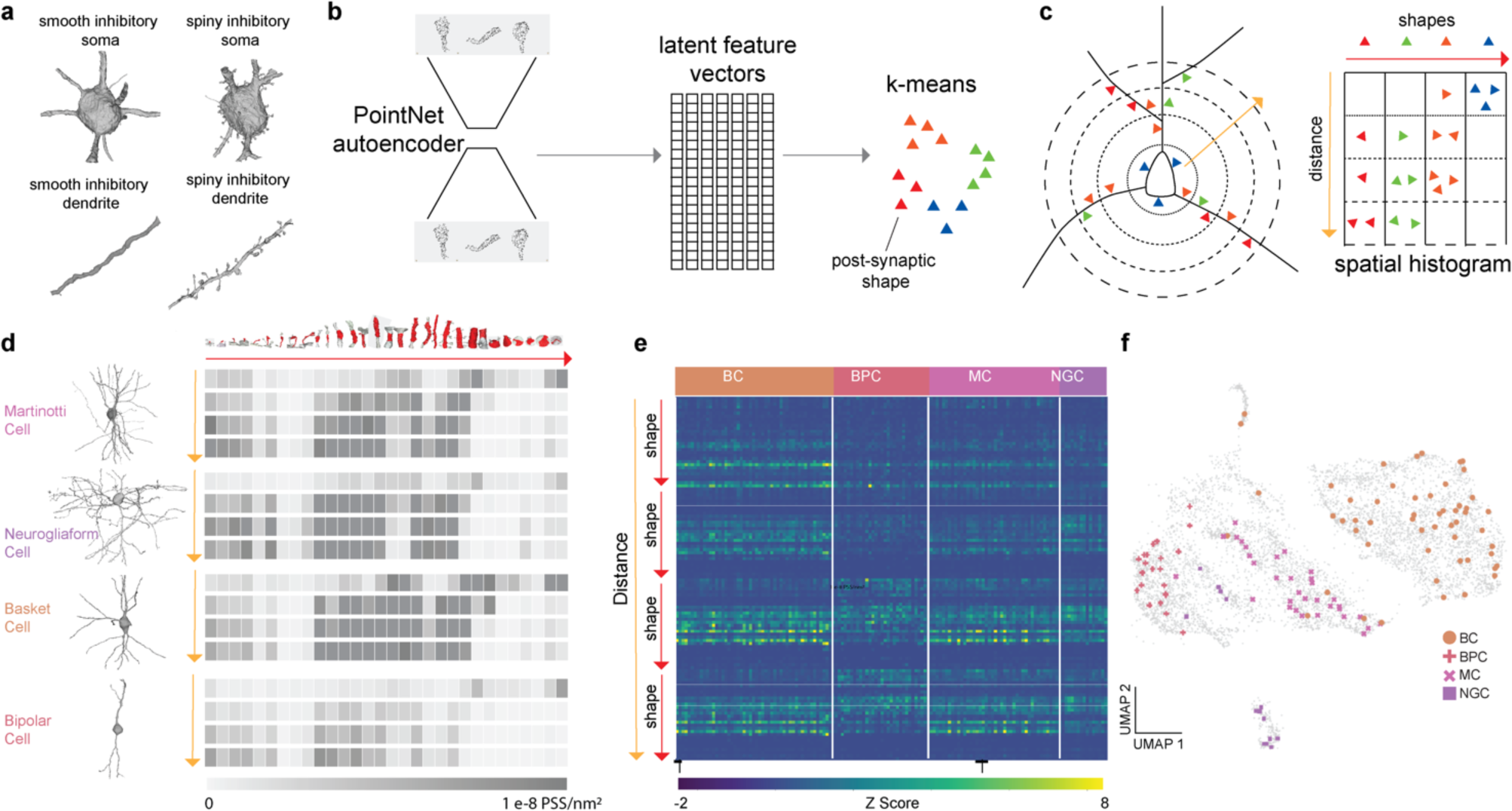
Post Synaptic Shape (PSS) Features. **a)** Inhibitory neurons elicit large variability in ultrastructural morphology. **b)** Procedure for building a PSS dictionary model. The set of shapes is used to train a PointNet autoencoder which learns a latent feature vector of a fixed size (1024). This autoencoder is then applied to all shapes in the dictionary to generate a set of latent feature vectors. K-means with K = 30 is applied to this to obtain a set of cluster centers for binning the shapes. **c)** For each cell, the PSS are binned by shape type and distance from the soma (4 bins) from 0 - 60 microns with 15 micron bin sizes. The resulting histogram is a 2D histogram shown above with the shapes in the × direction and distances in the y direction. **d)** Examples of 60 micron cutouts of the 4 predicted inhibitory subclasses with their spatial histograms shown as heatmaps. The top row shows the shape of the cluster center of each of the 30 clusters. In each heat map, darker boxes indicate higher values. **e)** Z-scored feature matrix representing the distance binned PSS features on the manually labeled inhibitory cells from the cortical column (n=143). Cells are organized by their annotated subclass. Dashed marks along the × axis denote segments of 100 cells **f)** 2D UMAP of all the inhibitory neurons (n=6,805) inferred after concatenating nucleus, soma and PSS features, cortical column cells in color and dataset wide inhibitory neurons in gray.

For this, we computationally segmented the compartment on the postsynaptic side of each synapse, which we refer to as the “Post-Synaptic Shape” (PSS).^27^ This shape, computed as a variable sized mesh, typically represented either a portion of the soma, the shaft of a dendrite, or a spiny protrusion, though it could also be onto an axon or axon-initial segment. In order to model the diversity of these shapes, we needed to be able to compare these shapes with each other computationally. We therefore trained a PointNet auto-encoder that allowed us to generate a fixed size representation of each shape (Fig. 4b) of dimensionality 1024.

We also wanted to measure the density of the distribution of shapes present in a cell. Consequently, we collected 236,000 PSSs from a variety of neurons and applied a 2D dimensional reduction in order to visualize their distribution. This resulted in a continuous latent space where PSS objects of similar morphological character were closer together (Extended Data Fig. 2). Since different cells can have different numbers of shapes (synapses), we now needed to develop a fixed size representation for a cell. For this, we used the similarity observation from the 2D latent space to develop a binning protocol for all the shapes. We used the full 1024 dimension features from the 236,000 PSSs and computed 30 cluster centers with K-means (Fig. 4b). Binning a set of shapes extracted on a cell using these cluster centers would therefore give us a 30 dimensional histogram. (The K-means algorithm randomly assigns the order of discretizations. Therefore, we also manually re-ordered the bin centers for visualization purposes from shapes representing small spines, to those representing longer spines, to dendritic shafts of different shapes, and finally somatic compartments.)

We also observed that the location of the PSS could give us extra cues to distinguish between cells. For example, while spiny protrusions were most often found on the dendrites of cells, some cells also had them on the soma (Fig. 4a, Extended Data Fig. 3). Therefore, we took a second step of summarizing a cell’s distribution of PSSs according to its shape and distance from the nucleus center. For distance binning we used four 15 micron bins between 0 to 60 microns from the soma (see Methods). Combining the shape and distance binning resulted in a 120 dimensional spatial shape histogram (Fig. 4c), that summarizes information about the spatial organization of dendritic shapes and synapse densities near the soma, similar to a multi-dimensional Sholl analysis.^28^ There were typically clear visual differences in the spatial histograms of different cell types (Fig. 4d-e, Extended Data Fig. 3). For example, a Martinotti cell had a greater density of synapses onto small protrusions on its proximal dendrites than the basket or bipolar cell, but similar numbers to the neurogliaform cell. However, the neurogliaform cell had very few synapses on its soma, whereas the Martinotti has many, both onto smaller protrusions and smoother compartments of its somatic compartment (Fig. 4d).

We then extracted these features on the vast majority of putative inhibitory neurons in the dataset (as predicted by perisomatic features, see methods). Appending these features to the soma features from the previous section and inspecting the UMAP (Fig. 4f) suggested that several inhibitory types that were not easily distinguishable without the PSS features were now more separable (subclass accuracy of 94%, Extended Data Table 1).

### Perisomatic Features Enable Dataset Wide Classification

In order to support these qualitative observations with quantitative methods and enable dataset wide classification, we trained a collection of classification models (support vector machines or multilayer perceptrons) on different feature sets: 1 - nucleus only, 2 - nucleus and soma, 3 - nucleus, soma and PSS (Extended Data Table 1). All classifiers were trained using the labels from the manually labeled cortical column (see methods for more detail). We developed a hierarchical model that used a cascade of classifiers to sort cells at increasingly finer distinctions and integrated the steps into a comprehensive model where individual cells are sequentially sorted down the hierarchical tree (Fig. 5a). We found an optimal combination of classifiers which predicted cell types labeled within the column with an overall accuracy of 91% (Fig. 5b, Extended Data Table 1), and importantly providing classifications for 88% of the cellular objects in the dataset (94,010/106,761 cells). To further validate this classification, we randomly sampled 100 cells from each subclass predicted by the hierarchical model and had human anatomical experts assess the labels (Extended Data Fig. 4). For many classes, the average classification in this validation was consistent with the accuracy found within the column. The lower validation accuracy in the inhibitory subclasses as well as 5P-ET and 5P-NP was likely related to the sparse sample sizes in the training data from the column. The largest single confusion between types here was between adjacent layers of similar pyramidal classes, where strict laminar boundaries separating manual classes is less confident. This demonstrates that these features are indeed useful for separating cell-types based on local somatic reconstructions of cortical cells, consistent with the structure of the low dimensional embedding (Fig. 5c). Furthermore, predictions of cell density and overall cell counts per subclass across the dataset (Extended Data Fig. 5) corroborate the sampling rates we would expect from previous studies.^29–32^ Importantly, this approach can easily be adapted to accommodate new cell-type labels derived from more detailed or expansive studies of the dataset, creating a scalable platform for extending labels derived on smaller numbers of cells to dataset wide coverage. For example, we have successfully trained models based on the unsupervised clustering labels of morphological and connectivity properties of the same column cells as described in Schneider-Mizell 2023 (Extended Data Fig. 6).

**Figure 5:**
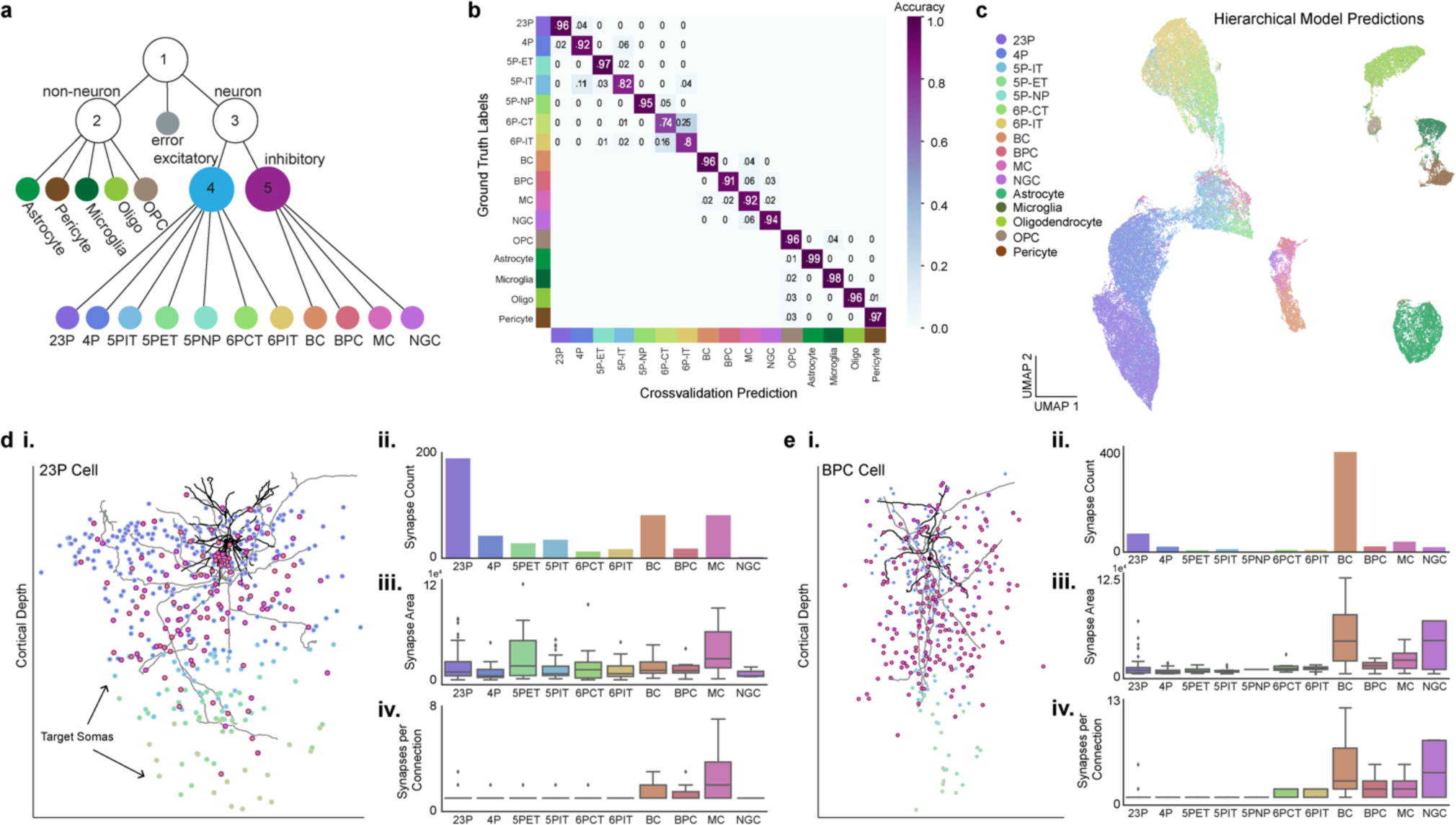
Hierarchical predictions enable dataset wide circuit analyses. **a)** Diagram of the hierarchical model framework used to predict neuronal and nonneuronal subclasses using a set of 5 classifiers. Nucleus and soma features alone were used for models 1-4. PSS features were added to predict inhibitory subclasses in model 5 (Extended Data Table 1). **b)** Confusion matrix of the cross validation performance for all cells within the manually labeled column. Note that classifiers for excitatory neurons, inhibitory neurons, and non-neurons were trained separately (Models 2,4,5 in panel a). The confusion rate between these classes can be seen in Extended Data Fig. 4. **c)** 2D UMAP embedding inferred from depth, nucleus, and soma features of all cells in the dataset colored by the hierarchical model predictions. **d) i)** 2D rendering of a representative 23P cell morphology, dendrite in black and axon in gray. Points represent the somatic position of all downstream target cells colored by the hierarchical model subclass prediction. **ii)** synapse count **iii)** total synapse area and **iv)** number of synapses per connection displayed by the model predicted subclasses illustrating the local targeting profile of this individual cell. **e)** Similar information as in d but for an inhibitory bipolar cell that is predicted to preferentially target basket cells. This unique population of bipolar cells has been further characterized in Schneider-Mizell 2023.

Dataset wide classifications enable a range of subsequent analyses. The typical axon of a well-proofread neuron has hundreds or thousands of postsynaptic targets.^3^ To quantify the cell-type specific connectivity of such cells, each of those targets should have a cell-type label. Doing so manually is a practical bottleneck in analyzing these data. With these predictions, scientists can easily analyze the most numerous postsynaptic targets, the weight of these synapses with respect to predicted synapse area, and the number of synapses between proofread cells and cell subclasses across thousands of synapses (Fig. 5d-e). For example, a given layer 2/3 pyramidal neuron made the most synapses onto other 23P neurons (Fig. 5d). However, when we looked at the total predicted synapse size, 5P-ET neurons receive some of the largest average synapses. On the other hand, some examples were more surprising than this 23P cell. For example, bipolar cells (which largely overlap with a VIP subclass) have been described as the only disinhibitory specialist interneuron class, and are described as making synapses primarily onto SST cells (which are thought to overlap largely with the Martinotti Cell definition used here).^33–36^ Although the dataset contains cells consistent with that view, a companion study on extensively proofread cells identified a collection of disinhibitory multipolar neurons which exhibit strong targeting preferences for basket cells.^16^ This unique connectivity profile is observed in the dataset wide classifications as well (Fig. 5e).

### Perisomatic Features Enable Efficient Search For Rare Cell Types

Studying the connectivity patterns of cell types requires identifying many example cells of a particular connectivity profile. With more than 70,000 neurons densely sampled across a millimeter scale there should be many examples of any individual cell type. However, locating those examples can be challenging for rare subclasses because of their infrequent appearance and the need for axonal proofreading in order to use connectivity to suggest their subclass.

Given that the major inhibitory neuron subclasses differ in their connectivity profiles, we already had some evidence that connectivity profile correlates with the perisomatic features we extracted (Fig. 4e), but we conjectured they could be useful for finding rarer types with highly specific connectivity patterns for which we did not yet have labels. One particularly interesting and well known rare cell type in mouse visual cortex is the chandelier cell, which exclusively synapses onto the axon initial segment (AIS) of excitatory neurons.^13–15,23,37,38^ We used a single proofread chandelier cell to see if we could facilitate finding other cells like it using the perisomatic features. We picked the top 20 nearest neighbors of the perisomatic feature space (Fig. 6a) and assessed what fraction of them were chandelier cells based on their connectivity profiles after cleaning them of false mergers and modest axonal extension (see methods). The chandelier cell’s connectivity profile is easy to recognize, both from its morphology where it makes vertical strings of synapses (Fig. 6b), and the unique targeting of synapses onto axon initial segment (AIS) of excitatory neurons. Because the AIS is usually located just below the soma of excitatory cells in the cortex, the angular distribution of synapses relative to somatic targets can be used as a spatial proxy for AIS targeting (Fig. 6c-d). A histogram of the angular distribution of synapses relative to the target soma (Fig. 6d-e), demonstrates that 16 of 20 the nearest neighbor cells have connectivities consistent with chandelier cells. In contrast, none of the 20 random interneurons we sampled from the inhibitory neurons in the dataset, or any of the 143 interneurons systematically sampled in the column were chandelier cells, reflecting a highly significant enrichment (p<0.00001 by Fisher exact test). Based on this success, we also tried to find more examples of cells with a less well known connectivity profile. We selected a classically undescribed but proofread interneuron which made the majority of its synapses in layer 5 onto 5P-NP neurons, despite those neurons being rare and with few input synapses (Fig. 6f).^39^ Picking the top 20 nearest neighbors of this cell, we found 13 cells which made at least 30% of their synaptic targets onto NP cells (based on our dataset wide classifier). This stands in contrast to the 0 of 20 random interneurons we sampled, or 2 of other cells of the other 163 systematically sampled in the column, again a highly significant enrichment (p<0.00001) (Fig. 6g). This application demonstrates how these features can be used to target rare cell types in the cortex.

**Figure 6:**
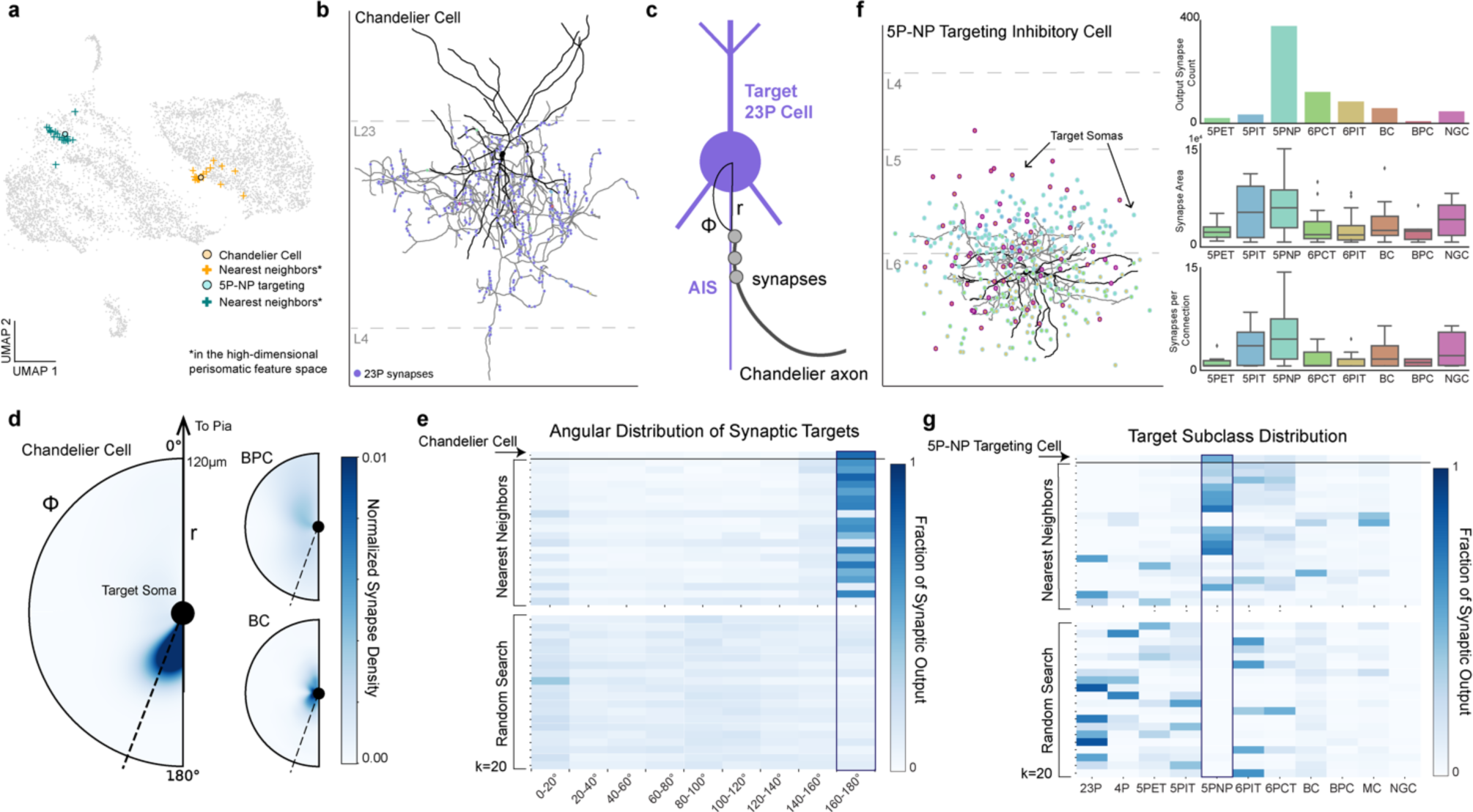
Perisomatic feature space enables more efficient search for unique cells. **a)** 2D UMAP embedding highlighting an example chandelier cell (orange dot) and an example 5P-NP targeting cell (blue dot) and their respective 20 nearest neighbors (+) in the perisomatic feature space. Note that UMAP non-linearly distorts feature space, so not all the nearest neighbors appear closest in the plot. **b)** Example proofread chandelier cell in layer 2/3 (dendrite in black, axon in gray). Output synapses marked along the axon and colored by the subclass prediction. Notice the characteristic vertical chains of synapses onto 23P cells. **c)** Chandelier cells are characterized by their preference to synapse onto the axon initial segment (AIS) of target cells.^13,14^ This can be quantified by measuring the angle between the target soma and the synapse (ɸ) and the distance from the soma (r). **d)** Heatmap illustrating the angle and distance distribution of the chandelier cell shown in B as well as two non Chandelier inhibitory examples. Note the spatial specificity of the chandelier cell targeting just below the target soma as compared to the other examples. Color notes the normalized synapse density for each cell. Synapses that had an angle >160° were considered onto the AIS of the target cell (shown by the dotted line). **e)** Angular distribution histogram of the chandelier cell (top row), the 20 nearest neighbors in the perisomatic feature space, and 20 random inhibitory cells. **f)** Example cell that preferentially targets the rare 5P-NP subclass (dendrite in black, axon in gray), points represent target cell soma locations colored by predicted subclass. Output synapse counts reflect strong preference to 5P-NP cells. **g)** Fraction of output connectivity onto neuronal subclasses of the 5P-NP targeting cell (top row), the 20 nearest neighbors in the perisomatic feature space, and 20 random inhibitory cells.

## Discussion

Our analysis of the perisomatic region of cells in the mouse visual cortex demonstrates that a surprising amount of cell-type information can be extracted from the somatic regions of brain cells. Our approach has already been used to characterize the connectivity of distinct types of layer 5 Martinotti cells,^17^ the inter-related connectivity motifs of layer 5 thick tufted cells (5P-ET) and the surrounding inhibitory sub-network,^18^ and to confirm the connectivity profiles of interneurons to cells outside the column.^16^ Future work in this dataset and others will likely leverage iterations of dataset wide cell classifiers to discover novel aspects of cell-type specific wiring of cortical circuits along with broader organizational principles of wiring. Other cell classification approaches have been applied to this dataset, including unsupervised clustering of morphological features, and supervised approaches based on morphological graphs.^40,41^ All these approaches have focused on smaller subsets of the data that contained higher quality or complete reconstructions, reducing their effective coverage in the datasets to less than half the cells. In contrast, by leveraging the perisomatic features our hierarchical model covers almost all cells and the majority of detected synapses in the dataset with a postsynaptic cell type. Furthermore, extracting these features from the 3D segmentation alone is both fast and computationally inexpensive, making this framework scalable and iterable across datasets.

The breadth of cells in large scale electron microscopy data makes it an attractive modality to study cell-types. Our approach provides an example of how computational methods are an important practical tool for efficiently directing study to small subsets of cells within large datasets. This is particularly clear within highly diverse and rare inhibitory cells (as shown in Fig. 5-6), but similar questions arise among glial sub-types. One such example is the difference between OPCs and premyelinating oligodendrocyte cells which are thought to be differentiated OPCs that are in transitional states to oligodendrocytes.^42^ The structural diversity of cells predicted as OPCs within the low dimensional embedding space (Fig. 5c) suggests that searching within the perisomatic feature space, as illustrated in Fig. 6, could be used to facilitate scientific discovery across brain cell types. More broadly, some of the features described here can be measured with other techniques, such as x-ray tomography or light microscopy, and can be used to distinguish cells into different subclasses in a manner similar to what has been presented here.

Many studies of anatomical diversity of cortical cells have focused on the diversity of dendritic and somatic morphologies, axonal projection patterns, and synaptic connectivity patterns.^24,43–45^ Fewer studies have focused on differences in the of somas,^46–48^ particularly quantitative studies of the 3d ultrastructure of the soma with large single cell sample sizes across all layers of cortex. Laminar differences in cell body size distributions are well known, and serve as the basis for cyto-architectural definitions of layers, which clearly correlates with shifts in cell-type distributions, particularly excitatory ones.^49^ For example, pyramidal Layer 5 ET projection neurons are characterized by their large somas. This likely reflects differing demands for gene expression and metabolic load.^50^ Also, 5P-NP neurons have been recognized before as having smaller rounder somas on average.^39,51^ Further, anecdotal descriptions of variations in nucleus in-folding have been reported, though only in two dimensions within a narrower range of types (https://pubmed.ncbi.nlm.nih.gov/3988983/). Intriguingly, differences in nucleus infoldings have been reported to be modulated by activity in some brain areas.^52^

One of the more striking features of our results is both the amount and distribution of perisomatic innervation varies across cell types. Although this has not been measured directly across a large number of cells and cell-types in cortex before, our results are consistent with other studies.^19,23^ In particular, in excitatory cells, somatic input has been noted to be dominated by inhibitory sources.^53^ Although this is less well characterized for inhibitory subtypes, some have larger fractions of excitatory input.^54,55^ In addition, different types of inhibitory axons show preference or avoidance of the somatic compartment.^53,54^ Because the total somatic synaptic density reflects an aggregate across all pre-synaptic types, such cell-type specific output patterns^54–56^ are likely related to and consistent with the variation we have observed across types in summed inputs.

There are a few limitations to this work that should be kept in mind when interpreting its results. First, our most detailed analysis has only been completed on one dataset which comes from a single animal. That said, some patterns are consistent with what was found in smaller published dataset from layer 2/3,^19,23,57^ and the basic patterns found in these features across mouse visual cortex are reproduced in a second smaller dataset (Extended Data Fig. 7). Our approach is not the final word in cell-type predictions in this dataset, or large scale EM in general, and there are a number of dimensions of potential improvement. First, cell-type labels will continue to evolve as more cells are classified by either human experts or quantitative methods with increasing specificity and sophistication. In particular, our validation results are consistent with there being a larger diversity of inhibitory cells than exist within the column and so expanding the number of classes and labels there could improve performance. However, we think the dataset wide framework we have presented here will continue to be valuable, as we still expect any new labels to only be available for a small subset of cells. It should be noted that while the soma and nucleus features are very fast, inexpensive, and scalable, the PSS features require significantly more computational resources. As such, we directed PSS feature analysis to the population of cells, inhibitory cells, that we believed warranted further differentiability. Second, this model does not make use of all the information present at the soma of neurons. For example, the detailed ultrastructure visible in the imagery is not fully utilized. Other methods have utilized the underlying imagery of cells to distinguish cell types, either through detection of more subcellular organelles like cilia or by using imagery more directly to define abstract embeddings.^56,58,59^ Such methods could augment the perisomatic features we have described here.

Beyond the somatic region, there are a large variety of studies have shown how local features visible in the ultrastructure contain information that encode information about cell types, including neurotransmitters of fly synapses, identity of neuromodulatory axons, or cutouts of local dendrite and axons.^56,60^ These results all support the view that large scale quantitative measurements of ultrastructure provides a rich basis for identifying cellular properties of cells. Focusing on the somatic region is particularly useful because volumes of all sizes encounter volume boundaries or limits in reconstruction accuracy and somas are singular locations of cells. We believe that the efficacy of this approach provides a roadmap for how to develop a scalable platform for leveraging local features of cells to infer cell-type classifications. Beyond neuroscience, this approach illustrates how large scale ultrastructural imaging of cells can facilitate study of highly diverse and rare cell populations if paired with appropriate quantitative analysis.

## Methods

### MICrONS Dataset

This dataset consists of a 1.4mm × 800µm × 800µm volumetric serial section EM dataset from mouse visual cortex of a male P87 mouse. The dataset covers all layers of cortex and spanning primary visual cortex and two higher visual areas. The dataset has been described in detail elsewhere.^3^ Briefly, two photon imaging was performed on the mouse, which was subsequently prepared for electron microscopy. The specimen was then sectioned and imaged using transmission electron microscopy.^1^ The images were then stitched, aligned, and processed through a deep learning segmentation algorithm, followed by manual proofreading.^1–3,16^

### Cortical Column

In this manuscript we leveraged proofreading and labels that were done as part of a separate study of a 100µm columnar region of primary visual cortex within the larger dataset.^16^ For clarity to the reader and completeness we are repeating some aspects of the methods that define that column here.

### Column Selection

The column borders were found by manually identifying a region in the primary visual cortex that was as far as possible from both the volume boundaries and the boundaries with higher order visual areas. A 100 × 100 µm box was placed on layer 2/3 and was extended along the y axis of the dataset. While analyzing the data, it was observed that deep layer neurons had apical dendrites that were not oriented along the most direct pia-to-white-matter direction, and thus adapted the definition of the column to accommodate these curved neuronal streamlines. Using a collection of layer 5 ET cells, points were placed along the apical dendrite to the cell body and then along the primary descending axon towards white matter. The slant angle was computed as two piecewise linear segments, one along the cortical depth to lower layer 5 where little slant was observed, and one along the direction defined by the vector averaged direction of the labeled axons.

Using these boundaries and nucleus centroids,^3^ all cells were identified inside the columnar volume. Coarse cell classes (excitatory, inhibitory, and non-neuronal) were assigned based on brief manual examination and rechecked by subsequent proofreading and confusion with early versions of the classifiers described here. To facilitate concurrent analysis and proofreading, all false merges were split connecting any column neurons to other cells (as defined by detected nuclei) before continuing with other work.

### Proofreading

Proofreading was performed primarily by five expert neuroanatomists using the PyChunkedGraph^57,61^ infrastructure and a modified version of Neuroglancer.^62^ Proofreading was aided by on-demand highlighting of branch points and tips on user-defined regions of a neuron based on rapid skeletonization (https://github.com/AllenInstitute/Guidebook). This approach quickly directed proofreader attention to potential false merges and locations for extension, as well as allowed a clear record of regions of an arbor that had been evaluated.

For dendrites, all branch points were checked for correctness and all tips to see if they could be extended. False merges of simple axon fragments onto dendrites were often not corrected in the raw data, since they could be computationally filtered for analysis after skeletonization (see below). Detached spine heads were not comprehensively proofread, and previous estimates place the rate of detachment at approximately 10-15%.

For inhibitory axons, axons were “cleaned” of false merges by looking at all branch points. Axonal tips were extended until either their biological completion or data ambiguities, particularly emphasizing all thick branches or tips that were well-suited to project to new laminar regions. For axons with many thousand synaptic outputs, some but not all tips were followed to completion once major branches were cleaned and established. For smaller neurons, particularly those with bipolar or multipolar morphology, most tips were extended to the point of completion or ambiguity. Axon proofreading time differed significantly by cell type not only because of differential total axon length, but axon thickness differences that resulted in differential quality of auto segmentations, with thicker axons being of higher initial quality. Typically, inhibitory axon cleaning and extension took 3-10 hours per neuron.Expert neuroanatomists further labeled excitatory and inhibitory neurons into subclasses. Layer definitions were based on considerations of both cell body density (in analogy with nuclear staining) supplemented by identifying kinks in the depth distribution of nucleus size near expected layer boundaries.

### Cell Labeling

For excitatory neurons, the categories used were: Layer 2/3-IT, Layer 4-IT, Layer 5-IT, Layer 5-ET, Layer 5-NP, Layer 6-IT, and Layer 6-CT cells. Layer 2/3 and upper Layer 4 cells were defined on the basis of dendritic morphology and cell body depth. Layer 5 cells were similarly defined by cell body depth, with projection subclasses distinguished by dendritic morphology following Gouwens, Sorenson, and Berg^9^ and classical descriptions of thick (ET) and thin-tufted (IT) cells. Layer 5 ET cells had thick apical dendrites, large cell bodies, numerous spines, a pronounced apical tuft, and deeper ET cells had many oblique dendrites. Layer 5 IT cells had more slender apical dendrites and smaller tufts, fewer spines, and fewer dendritic branches overall. Layer 5 NP cells corresponded to the “Spiny 10” subclass described in Gouwens, Sorenson, and Berg; these cells had few basal dendritic branches, each very long and with few spines or intermediate branch points. Layer 6 neurons were defined by cell body depth, but only some cells were able to be labeled as IT or CT by human experts. Layer 6 pyramidal cells with stellate dendritic morphology, inverted apical dendrites, or wide dendritic arbors were classified as IT cells. Layer 6 pyramidal cells with small and narrow basal dendrites, an apical dendrite ascending to Layer 4 or Layer 1, and a myelinated primary axon projecting into white matter were labeled as CT cells.

Basket cells were recognized as cells which made more than 20% of their synaptic inputs onto the soma or proximal dendrites of cells. Neurogliaform cells were recognized by having a low density of output synapses, and boutons that had often had synaptic vesicles but no post-synaptic structures. Bipolar cells were labeled by having only 2 or 3 primary dendrites, and primarily making synapses with other inhibitory neurons. Note, the Martinotti/non-Martinotti subclass label was given to cells that have previously been described in the literature to primarily target the distal dendrites of excitatory neurons without exhibiting hallmark features of bi-polar or neurogliaform cells.

Due to high levels of proofreading in the column, there were very few errors thus the training set was augmented with manually labeled errors from the entire dataset.

### Proofreading and Truncation Analysis

For every proofread cell in the cortical column (described above) we compared the cellular volume of the initial reconstruction from the automated segmentation to the cleaned and completed reconstruction. To measure the precision connectivity for each cell we noted the number of synapses that got removed with proofreading, the number of synapses that were added, and the number of synapses that were maintained with each cell before and after proofreading.

To estimate the likelihood of truncation, we measured the distribution of dendritic extents from the proofread column cells. For each cell we measured the radial distance of each input synapse from the cell’s soma. The radial extent of a given cell was considered the distance of the 97th percentile input synapse. From this distribution we used the median value of 121 microns as a threshold for dendritic truncation, although closer to 250 microns would be required to guarantee no truncation for any cell. For the rest of the cells in the dataset, we measured the distance of the soma from the volume borders in × and z. The overlap in these distributions relates to the probability of truncation, leading to our conclusion that roughly one third of the cells have some degree of dendritic truncation.

### Generating Nucleus and Soma Features

We analyzed nuclei using the results of a deep neural network segmentation,^3^ extracted the mesh using marching cubes and obtained the largest component of the detected mesh. Nuclear features were then extracted on the remaining meshes. These features included, nucleus volume, nucleus area, the area to volume ratio, nucleus surface area within an infolding, the fraction of the total surface area within an infolding, and cortical depth (measured as the distance from the pial surface). Nucleus fold features were extracted by creating a shrink wrapped^47^ mesh for each nucleus mesh. We then calculated the distance of each vertex on the nucleus mesh from the shrink-wrapped mesh. After visual inspection of cells across all the reported subclasses, any vertex further than 150 nm was considered within an infolding.

For each nucleus detection the somatic compartment was identified as the ID in the segmentation which surrounded >80% of the nucleus. Somatic segmentations (downloaded at 64×64×40 nm resolution) went through a heuristic cleaning procedure to remove missing slices of data and incorrectly merged fragments. Since each soma was matched to its corresponding nucleus, 15 microns surrounding the nucleus’ center of mass was cut out from the dense segmentation and converted into a binary mask. 15 microns was chosen due to the high quality of the segmentation (Fig. 2a) and it was large enough to encompass the entire soma of all cells from the smallest glial cell to the largest 5P-ET neuron. Binary dilation by 5 voxels in 3d was performed, followed by filling of all holes, and then binary erosion of 3 voxels. The resulting binary mask was meshed using marching cubes and connected component analysis was run on the result. 5 voxels was deemed an appropriate dilation to remove merged fragments without creating additional holes in the mesh. The largest connected component mesh was retained, and any disconnected components were dropped. Somatic features were extracted for all nuclear detections that were not cut off by the volume boundary (see Filtering procedure). These somatic features included soma area, soma volume, the area to volume ratio, the number of synapses on the somatic cutout, and the soma synapse density. Using both the somatic and nucleus meshes, we calculated the ratio between the nucleus volume and soma volume and the offset between the two, measured as the euclidean distance between nuclear center of mass and soma center of mass.

### Filtering procedure

There were 133,580 nuclear detections in the dataset and the filtering procedure consisted of three steps. Firstly, any detected objects less than 25µm^3^ were filtered out as errors as these largely consisted of small fragments of nucleoli. Second, after identifying the segment IDs within a 15μm bounding box around each nucleus, if over 20% of these IDs corresponded to error ID 0, they were filtered out. The majority of these error cases were cells close to the volume border or areas in the volume with higher segmentation errors such as those near blood vessels. Thirdly, cells that were predicted as errors based on the object classifier of the hierarchical model described below (Fig. 5a) were also removed from analysis. This resulted in a final set of 94,010 cells, neuronal and nonneuronal.

### Feature normalization

Due to differences in section thickness during sample preparation, we noticed abrupt shifts in nucleus and soma size features along the sectioning axis (Z plane). This presumably is due to changes in section thickness across the dataset. To account for these abrupt and systematic shifts we binned the entire dataset by the longest length scale for which there didn’t appear to be systematic shifts in the distribution in the z plane (800 nm) and normalized each feature value by the average within each Z bin.

For 2D UMAP embeddings and training of the classifiers it was important to place all features in approximately similar scales. For this reason, we independently Z-scored each feature across all cells and used that as the input for classifier training as well as the UMAP embeddings in Fig. 3-6.

### Generating PSS Features

Around each synapse, we extracted a 3500 nm region to obtain the synapse region mesh. We experimented with region cutouts between 1000 to 5000 nm, however smaller cutouts led to ambiguities in the main shaft identification and thereby produced errors in the subsequent skeletonization. At 3500 nm the skeletons were more stable and segmenting as expected. This mesh was then segmented using the CGAL surface segmentation algorithm^63^ which splits regions based on differences in thickness. We adapted our previously developed method^27^ to identify the PSS region by using a local skeleton calculated from the synapse region mesh, rather than a precomputed whole cell mesh. This allowed us to adapt this method for cells in the dataset without the need for proofreading.

Given a cell for which all PSS have been extracted within a 60 micron radius from the nucleus center, the objective was to build a descriptor that encapsulates the various properties of the PSS. Initially we extracted PSS from within 120 micron radius. However, upon inspection of the normalized histograms and the 2D UMAP embedding space, the additional radial bins did not increase our differentiability and did increase truncation effects near the dataset thus we reduced the radius to 60 microns. In particular, we aim to capture two of these properties: the type of shape of the PSS and the distance of the PSS from the soma. For the shape, a dictionary of all shape types is built using the dictionary dataset from.^27^ These shapes were rotationally normalized and used to train a pointnet autoencoder^64^ to learn a latent representation of size 1024. The high dimensional latent space spanning all these shapes is a continuous space (Extended Data Fig. 3) which was used to generate a Bag of Words model^30^ for the shapes. To ensure we were sampling the entire embedding space, we performed K-means clustering with K=30 to estimate cluster centers. The top row of Fig. 4c shows the shape in the dictionary that is closest to each of these cluster centers. For distance binning, we split the 60 micron radius around the nucleus center into four 15 micron radial bins (Fig. 4c). All PSS were then binned according to their shape and distance properties to generate a histogram of counts. This histogram was Z-scored and then added to the rest of the features as input to classifiers and the UMAP embedding Fig. 4 and Fig. 6.

### Hierarchical Model Training and Validation

#### Hierarchical Framework

We defined an object as the segmentation associated with a predicted nucleus^3^ from which nucleus, soma, and PSS features could be extracted. A hierarchical framework was designed to predict the cell type of any such object (Fig. 5c). To begin, there were 106,761 nuclear segmentations that passed the first two filters described above (see filtering procedure). The first level in the hierarchy predicted whether an object was a neuron (72,158), nonneuron (21,856), or an error (12,751). All objects predicted as errors were excluded from all subsequent analyses except for the hierarchical model evaluation. Nonneuronal cells were then classified as one of the following: astrocyte (7,850), microglia (2,638), oligodendrocyte (7,020), oligodendrocyte precursor cells (OPC) (1,703), or pericyte (2,645). For neurons, cells were predicted as either excitatory (64,195) or inhibitory (7,963) followed by a separate subclass classifier for each class type. Excitatory subclasses: Layer 2/3 pyramidal (19,735), Layer 4 pyramidal (14,777), Layer 5 IT (7,949), Layer 5 ET (2,215), Layer 5 near projecting (NP) pyramidal (970), Layer 6 IT (11,734), Layer 6 CT pyramidal (6,815). After extracting PSS features from all predicted inhibitory neurons, a subset of neurons (n=1,158) that were actually excitatory clearly separated from the rest of the cells in the perisomatic feature space (with PSS features). This was expected due to known differences in proximal dendrite morphology between inhibitory and excitatory neurons. These neurons were then passed through the excitatory neuron classifier and labeled as excitatory for all subsequent analyses with a final set of 6,805 inhibitory cells with the following subclass counts: Basket cells (3,239), Bipolar cells (997), Martinotti/non-Martinotti cells (1,992), and Neurogliaform cells (571).

#### Training

Soma and nucleus features were extracted from the 3D mesh of all objects and PSS features were extracted for all neurons predicted as inhibitory. For each level of the hierarchy, multiple classifiers were trained using either nucleus only, nucleus and soma features, or nucleus, soma, and PSS features. Within each level of the hierarchy, classifiers were trained using the cells and labels from the manually annotated cortical column. Due to the sparsity of some of the cell classes, we augmented the training set in the following ways: 470 errors were added from within and around the column for the object model, 11 proofread 5P-NP cells and 250 proofread 5P-ET cells were added to train the excitatory subclass model.

For each classifier, model type was chosen using a randomized grid search for the following models: Support Vector Machine SVM with a linear kernel, SVM with a radial basis function kernel, Nearest Neighbors, Random Forest Classifier, Decision Tree and Neural Network. For each type, 50 models were trained with varying parameters and the top performing model was chosen. Individual models were further optimized using 10-fold cross validation evaluated based on accuracy and F1 score (a measure for precision and recall). Training, and test examples were held consistent across models for direct performance comparison within each level.

#### Model Performance and Validation

The hierarchical model was defined as the sequential combination of the best performing classifiers at each level. To see the performance of the all different feature sets at each level of the hierarchy please see Extended Data Table 1. The overall performance of the hierarchical model was measured with a test set that involved manual inspection of 100 examples of each of the neuronal and nonneuronal subclasses as well as errors. This resulted in a test set of 1700 cells. Cross validation and test performance for the hierarchical model are reported below (Extended Data Fig.4). Note that all scores reported are the weighted accuracy based on the sampling rate of each class within the column.

The top level of the hierarchy (the object model), distinguished neurons from non-neurons as well as erroneous detections. The cross validated accuracy score on the column was 96% with a test score of 97%. The second level of the model simply distinguished excitatory from inhibitory neurons. Here, the column cross validated accuracy score was 94% and the test set was 93%. Overall, across all subclasses, the hierarchical model on the column had a cross validated accuracy of 91% and a dataset wide test set accuracy of 82%.

### Chandelier Cell Identification

Chandelier cells are characterized by their unique axo-axonal synapses onto the AIS of target pyramidal cells. As there were no chandelier cells within the densely reconstructed column, we sought to test if the perisomatic feature space would facilitate an enriched dataset wide search for these cells. After identifying and proofreading a chandelier cell, we selected the top 20 nearest neighbors by euclidean distance using a KDTree search of the perisomatic feature space (nucleus, soma, and PSS features) after z-score normalization of each feature across cells. We also selected 20 random cells from the predicted inhibitory neurons. For each of these 40 cells, we proofread the reconstructions to ensure that there were no extraneous neurites attached, and extended the axon until there were at minimum 100 output synapses. On average the 20 nearest neighbors had 590 output synapses attached and the random cells had 809 synapses attached.

To quantify whether a given cell was a chandelier or not, we measured the angle (ɸ) and the distance (r) between every output synapse and the soma of the postsynaptic cell (Fig. 6c). A synapse with an angle value of 0° would be considered straight above the target soma whereas an angle of 180° would be right below. Due to variations in axon directionality with respect to the pial surface, we determined that synapses with angle values between 160-180° and within 60 microns of the soma were considered on the AIS of the target soma. In fact, because the specificity of chandelier targeting is so high, the density of synapse angle distributions alone was enough to identify other chandelier cells (Fig. 6e). Upon inspection of the proofread 20 nearest neighbors, we determined that cells with over 40% of their synapses within 160-180° were chandelier cells. The average normalized density for the identified cells was 62% as compared to 8% for the non chandelier cells.

### Inhibitory Neuron Output Targeting

After characterizing a single 5P-NP targeting cell, we applied a similar strategy to the one above to search for more neurons in the dataset that had a similar connectivity pattern. We selected the top 20 nearest neighbors by euclidean distance in the perisomatic feature space using KDTree search. These cells were proofread to remove false mergers and extend the axon to include at minimum 100 synapses. It should be noted that there were 5 cells where the axons could not be extended due to volume boundaries or segmentation errors so they were replaced with the 5 nearest cells. On average the 20 nearest neighbors had 448 synapses attached.

To quantify whether a cell preferentially targeted 5P-NP neurons, we measured the fraction of total output that targeted different predicted subclasses. Cells that output over 30% of their synapses onto 5P-NP cells were considered to have this rare connectivity preference.

### Predicted Subclass Densities

To measure the predicted cell densities per subclass across the MICrONS dataset, we divided the dataset into 50µm^2^ bins in the XZ plane. Within each bin we calculated the number of cells for each subclass and scaled that to a mm^2^ to facilitate direct comparisons to reported densities in the literature.

### Dataset 2

The second dataset covers a millimeter square cross-sectional area, and 50 microns of depth within the primary visual cortex of a P49 male mouse.^19,23,57^ The largest available segmentation spans Layer 2/3 of the cortex through to Layer 6. After applying the nuclear detection model^19^ and filtering out all nuclear objects below 25µm^3^ and cells that were cut off by the volume border (see Filtering procedure above), 1,944 cells were used for the analysis. Class type of each cell was labeled manually and used as ground truth. Due to thinness of the volume, much of the distal cell morphologies were cut off and thus subclass type labeling was not possible. Nuclear and somatic mesh cleaning as well as feature extraction and normalization followed the same procedures outlined above.

All datasets described in the manuscript are publicly available at https://microns-explorer.org/ and https://bossdb.org/

## Contributions

Analyzed Data: LE, SS, FC

Developed Nucleus Model: SM, GM, LE, FC Cell

Typing: AB, NDC, CSM, JB

Paper Writing: LE, SS, FC

Dataset + segmentation generation: collaboration Proofreading: collaboration Analysis infrastructure: collaboration

## Support

LE was supported by RF1 MH117808

## Supplemental Information

**Extended Data Table 1:** Cross validation accuracy scores for individual classifiers at each level of the hierarchical model with differing input features. Each row corresponds to the corresponding numbers in the diagram in Fig. 5A. All training examples were held consistent between features sets for appropriate model comparisons. Classifiers with the highest accuracy score at each level were included in the hierarchical model (shown in bold). The overall hierarchical model performance on the column and the dataset wide validation set (see methods) is reported at the bottom.

|  | Nucleus<br>Column Cross-Validation<br>Accuracy | Soma + Nucleus<br>Column Cross-Validation<br>Accuracy | PSS + Soma + Nucleus<br>Column Cross-Validation<br>Accuracy |
| --- | --- | --- | --- |
| 1. Object Class | 0.90 | **0.96** | — |
| 2. Nonneuron Class | 0.94 | **0.98** | — |
| 3. Neuron Class | 0.95 | **0.99** | — |
| 4. Excitatory Subclass | 0.88 | **0.90** | — |
| 5. Inhibitory Subclass | 0.63 | 0.90 | **0.94** |

**Extended Data Table 1: Cross validation accuracy scores for individual classifiers at each level of the hierarchical model with differing input features.**
|  | Column Cross-Validation<br>Accuracy | Dataset Wide Validation<br>Accuracy |
| --- | --- | --- |
| **Hierarchical Model** | **0.91** | **0.82** |

**Extended Data Figure 1:**
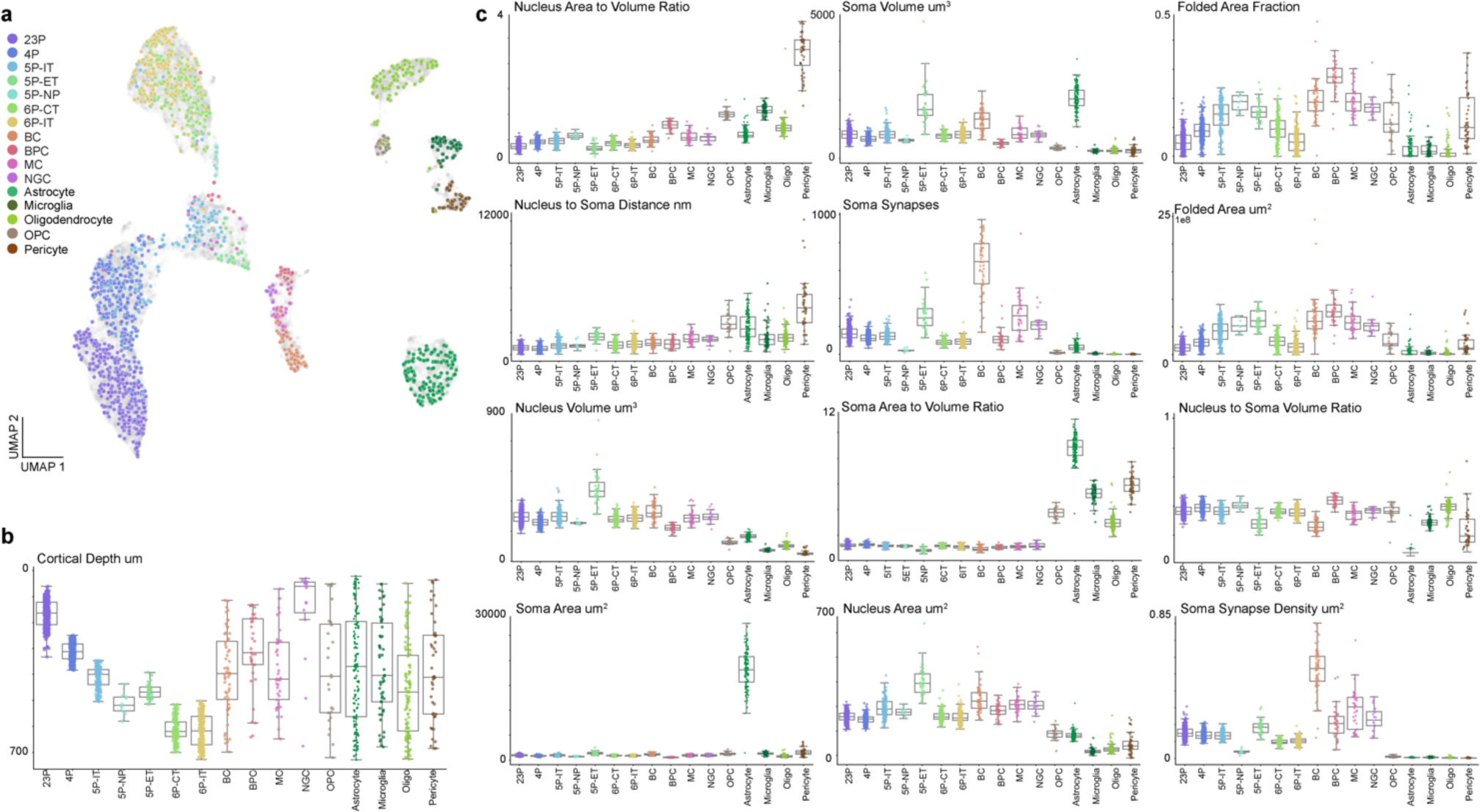
Neuronal and nonneuronal subclass distribution of individual soma and nucleus features. a) 2D UMAP embedding of all neuronal and nonneuronal cells inferred from somatic features, nuclear features and cortical depth. Manually labeled cellular subclasses are represented in color (1,619) and unlabeled examples in light gray (n=92,391). b) Distribution and variation of cortical depth of all cells from the human labeled column dataset. c) Distribution and variation of nucleus and somatic features of all cells from the column dataset. For all plots, mean and variance each subclass represented by the boxplots while individual cells are noted in the overlaid swarm plots. Color denotes human assigned subclass labels.

**Extended Data Figure 2:**
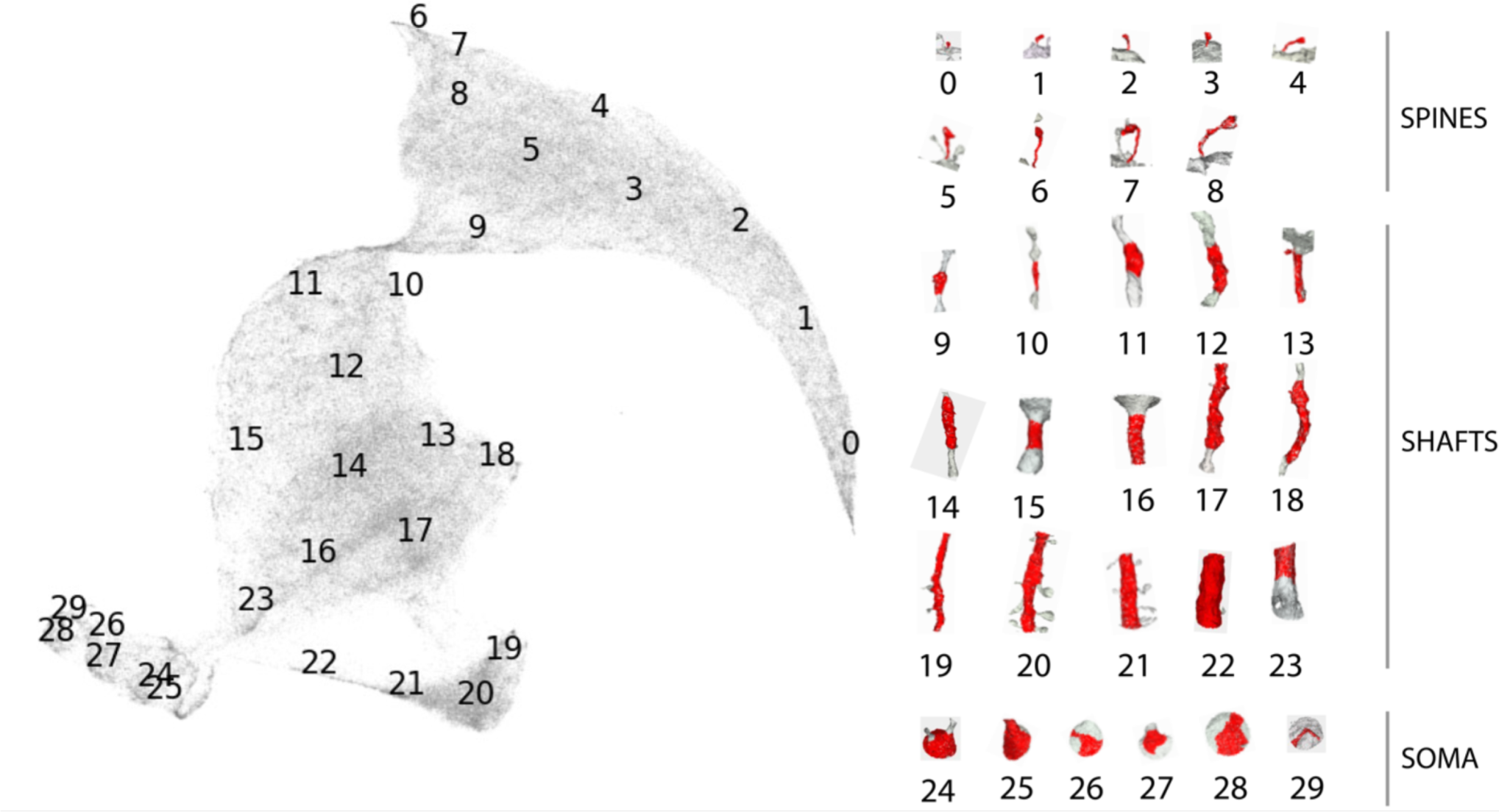
PSS embedding space organized by post-synaptic ultrastructural morphologies. 2D UMAP embedding of all shapes in the PSS Dictionary. The numbers indicate the bin centers mapped in this 2D space and the corresponding PSS meshes on the right show the shape associated with each bin center. Bins 1-8 range in spine shapes, Bins 9-23 are shaft shapes and Bins 24-29 are soma shapes.

**Extended Data Figure 3:**
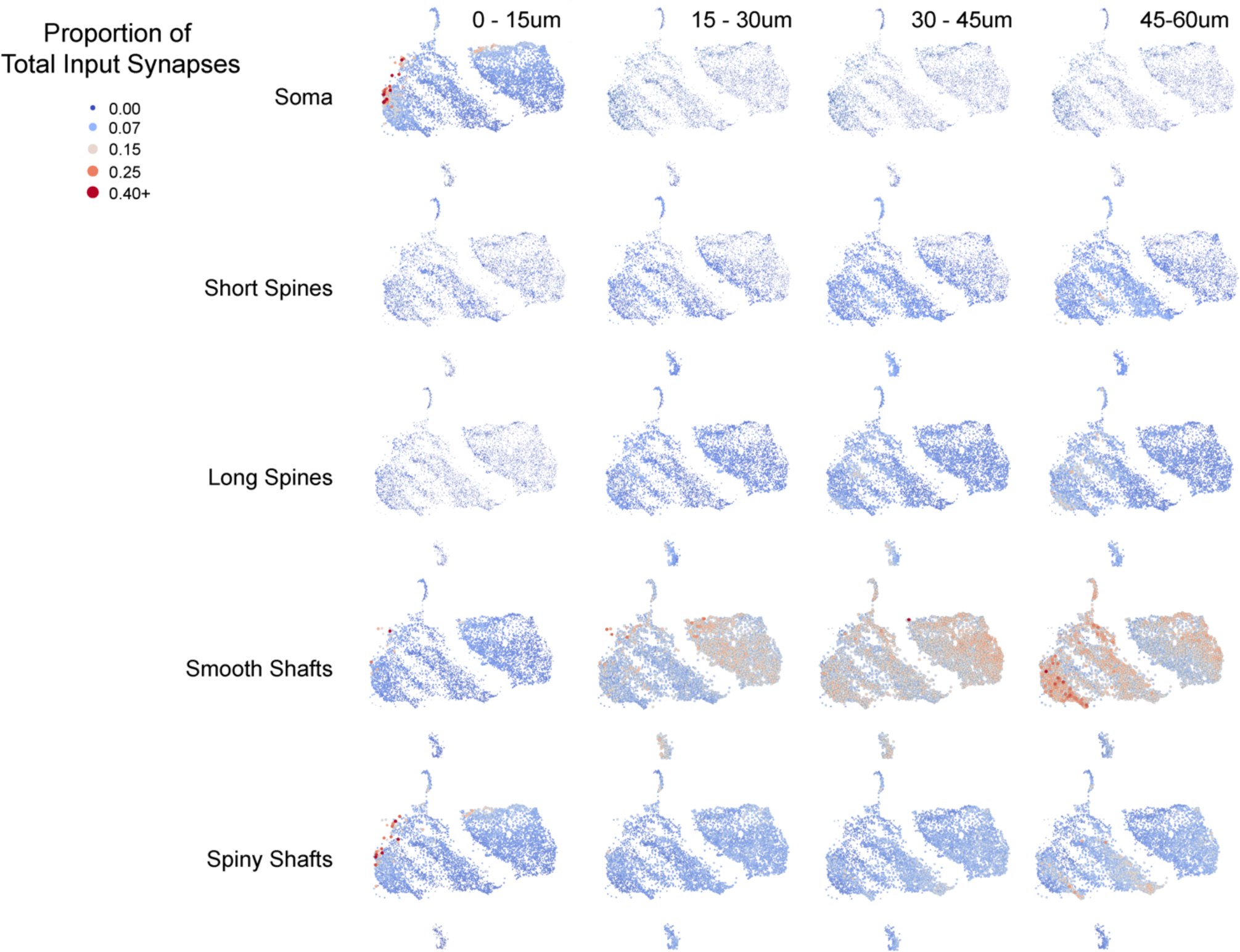
Inhibitory neuron subclasses exhibit spatial patterns to PSS distributions. The UMAP embedding of all the perisomatic features, including PSS features, across all inhibitory cells, colored with respect to what fraction of that cell’s input (within the 60µm cutout) comes from what PSS/distance bin. PSS shape bins were simplified from 29 bins to 5 broad categories to simplify the visualization (bins 0-4: short spines, 5-8: long-spines, 9-18+23: smooth shafts, 19-22: spiny shafts, 24-29: soma). This visualization gives insight into how different cells in different parts of this embedding space receive varying amounts of input onto different shapes within different spatial zones of the perisomatic area. Cells on the far left hand side of the embedding, where in general bipolar type neurons were found, have larger fractions of their inputs near the soma, including dendritic shafts which are more irregular in shape (“spiny shafts”), and smooth shaft inputs farther away where the dendrites begin to elaborate. Basket cells on the right hand side of the side of the embedding are dominated by somatic inputs and smooth shaft inputs which are more evenly distributed spatially. The island at the bottom that is dominated by neurogliaform cells is characterized by having relatively fewer somatic inputs, but an increasing amount of shaft and spiny input at distal dendrites.

**Extended Data Figure 4:**
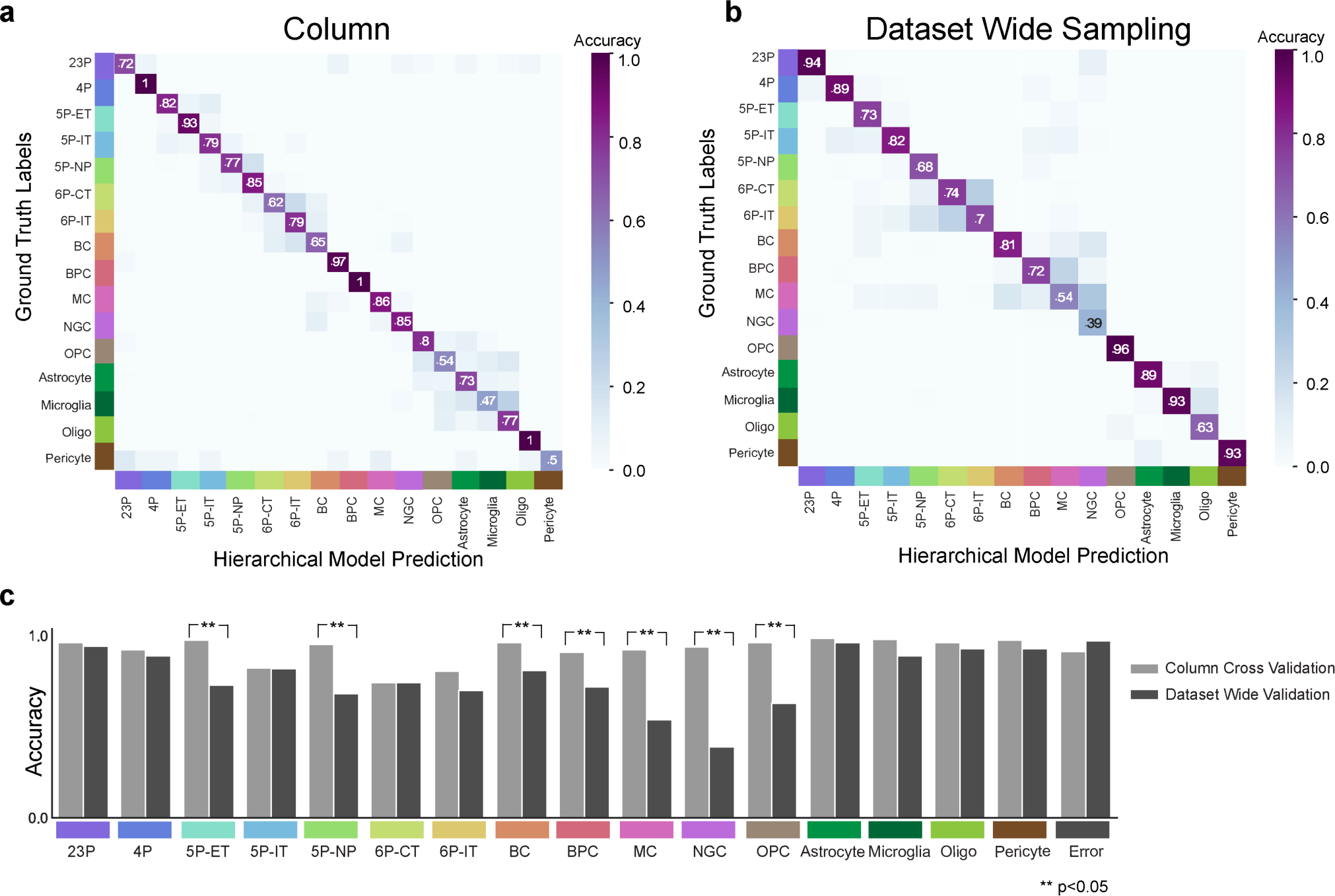
Classifier validation. **a)** Confusion matrix of hierarchical model performance for all cells within the manually labeled column after training. **b**) Confusion matrix of hierarchical model performance on a dataset wide sample of 100 cell predictions from each subclass. **c**) Comparison of column cross validation vs. dataset wide model performance, asterisk notes significance by Fisher Exact Test.

**Extended Data Figure 5:**
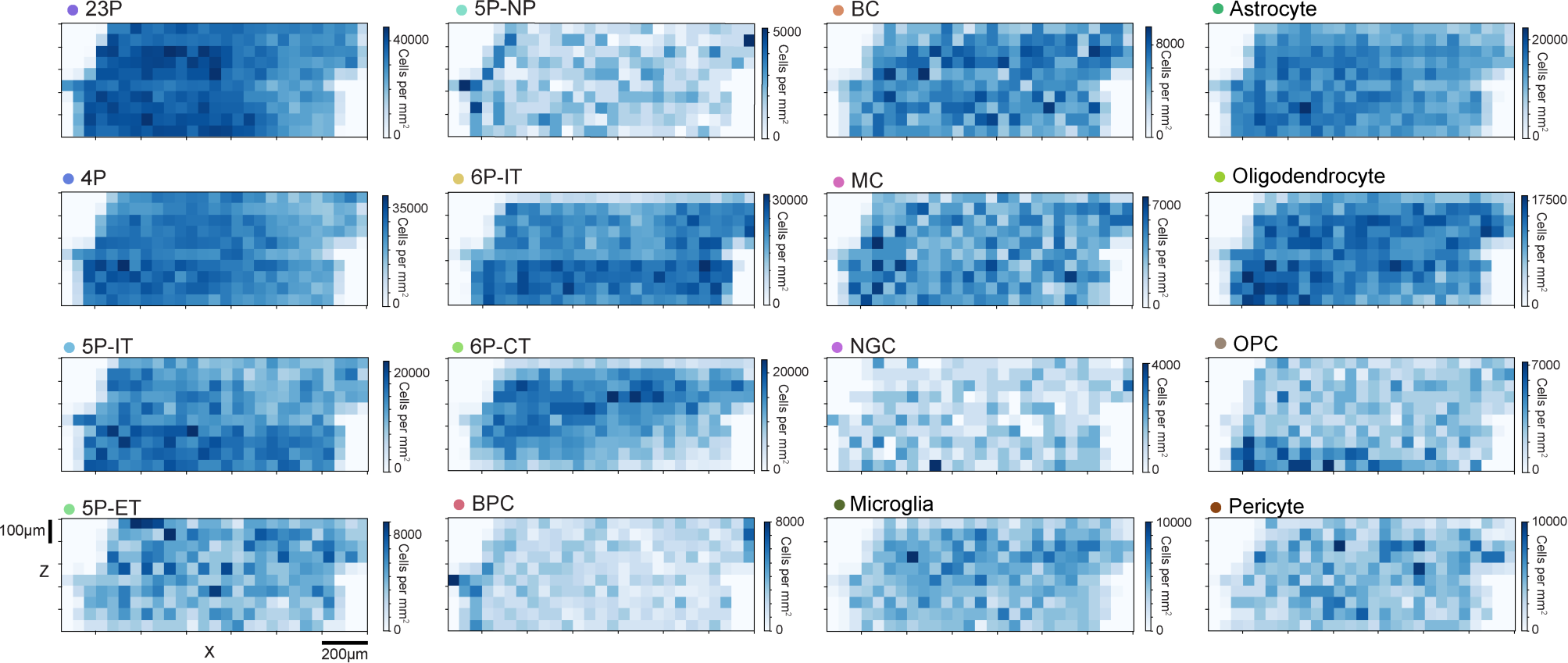
Cell densities across the dataset by cellular subclass. Predicted cell densities per mm^2^ for each subclass across the entire dataset in the XZ plane. Each square represents 50 micron^2^ and color denotes the density scaled per mm^2^. Note due to the approximate 1 mm depth of cortex, these values are also roughly densities per mm3. They roughly agree with densities of cells estimated from light microscopy stereology of subclasses,^29^ usually utilizing histochemical markers or genetic tools. Unfortunately for some subclasses, there is not a 1-1 to alignment between the definitions of types in this study and the usual molecular markers used in those studies, as molecular markers are not directly measurable in this electron microscopy volume.

**Extended Data Figure 6:**
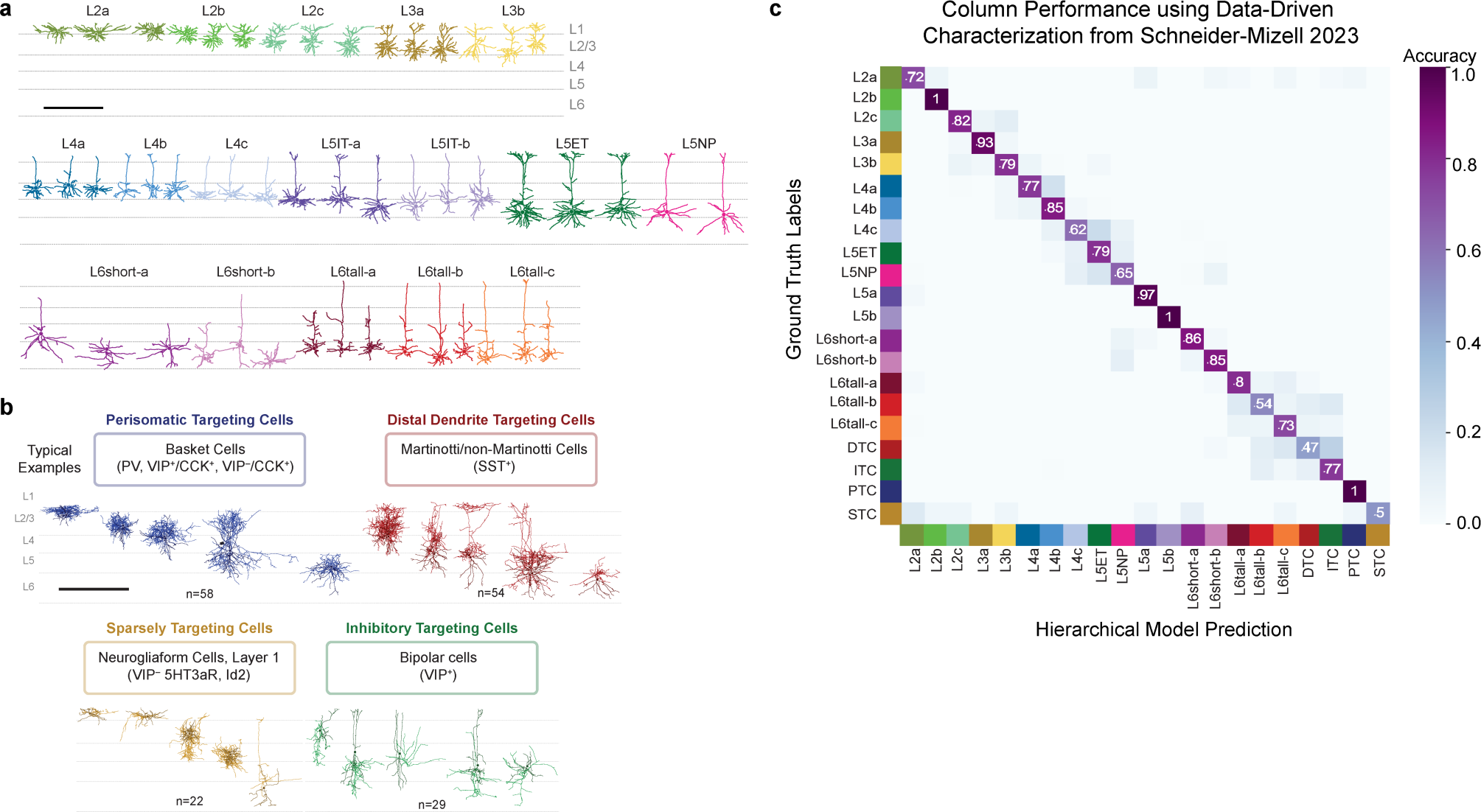
Perisomatic feature based classification utilized with different cell-type labels. **a)** Alternative excitatory subclass labels in the column from Schneider-Mizell et al 2023, based on unsupervised clustering of dendritic and synaptic features rather than manual human expert calls. Labels on the clusters were inferred based on the overlap with expert labels and cortical depth, with finer distinctions added when necessary (i.e. L4a,L4b,L4c). **b)** Alternative inhibitory subclass labels from Schneider-Mizell et al. 2023 in the column based on unsupervised clustering of their output connectivity statistics. These subclasses (Perisomatic targeting, Distal Targeting, Sparsely Targeting and Inhibitory Targeting) likely largely but not completely align with broad molecular distinctions made amongst inhibitory cells, based on reviews of the literature where molecular and output connectivity has been measured in the same cells. **c)** A confusion matrix of a hierarchical model retrained to utilize these subclass labels for excitatory neurons vs inhibitory neurons rather than human expert labels. Cross validation performance on the excitatory (67%) and inhibitory (85%) subclass models was lower than the expert labels, due primarily to the fine grained distinctions made amongst layer 4 and 6 types. The confusion matrix shown here is the output of the final model trained on all samples from the column.

**Extended Data Figure 7:**
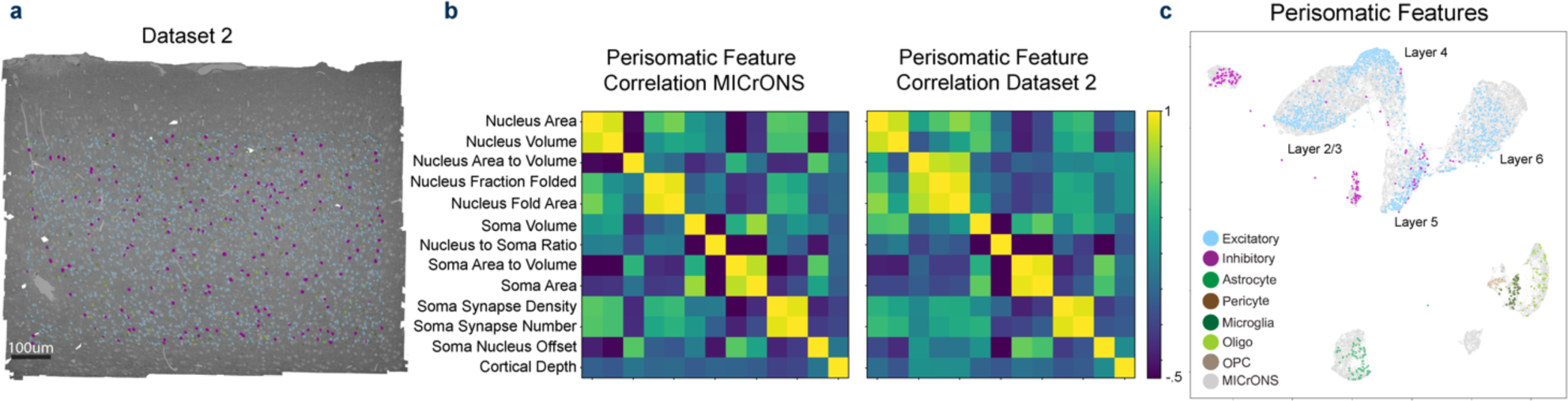
Basic perisomatic feature patterns maintained across a second dataset from a different animal. **a)** A cutout of a second dataset, which covers layer 2/3 to 6 of cortex, but is only 50µm thick. Somas contained within this volume (n=1,944) were analyzed in a manner identical to the larger dataset and soma, nucleus and PSS features were extracted. Excitatory nuclei highlighted in light blue and inhibitory nuclei in magenta. **b)** Analysis of the feature to feature correlations shows similar correlation structure between the two datasets. **c)** A joint UMAP of the perisomatic features with the MICrONS dataset data shown in gray, and the smaller dataset covered by manually identified cell classes overlaid. In general, the same overall patterns and degree of separation amongst layers and cell classes was observed. Note: pericytes were manually excluded from this dataset due to the lower quality of nucleus and somatic segmentations. Extensive detailed subclass cell type validation is not possible in this dataset due to the truncation of axons and dendrites.

## Notes

https://www.microns-explorer.org/

### Competing Interest Statement

The authors have declared no competing interest.

### Summary of Updates

Figure 1-3 updated to clarify the feature discriminability. Figure 6 updated to clarify the application of the method. Supplemental files updated.

